# IRF8 Governs Tumor-Associated Macrophage Control of T Cell Exhaustion

**DOI:** 10.1101/2020.03.12.989731

**Authors:** Briana G. Nixon, Fengshen Kuo, Ming Liu, Kristelle Capistrano, Mytrang Do, Ruth A. Franklin, Xiaodi Wu, Emily R. Kansler, Raghvendra M. Srivastava, Tanaya A. Purohit, Alejandro Sanchez, Lynda Vuong, Chirag Krishna, Herbert C. Morse, James J. Hsieh, Timothy A. Chan, Kenneth M. Murphy, James J. Moon, A. Ari Hakimi, Ming O. Li

**Affiliations:** Immunology Program, Sloan Kettering Institute, Memorial Sloan Kettering Cancer Center, New York, NY, USA 10065; Immunology and Microbial Pathogenesis Program, Weill Cornell Graduate School of Medical Sciences, Cornell University, New York, NY, USA 10065; Immunogenomics & Precision Oncology Platform (IPOP), Memorial Sloan Kettering Cancer Center, New York, NY, USA 10065; Department of Pathology and Immunology, Washington University in St. Louis, School of Medicine, St. Louis, MO, USA, 63110; Howard Hughes Medical Institute, Washington University in St. Louis, School of Medicine, St Louis, MO, USA, 63110; Urology Service, Department of Surgery, Memorial Sloan Kettering Cancer Center New York, NY USA, 10065; Computational and Systems Biology Program, Memorial Sloan Kettering Cancer Center New York, NY USA, 10065; Virology and Cellular Immunology Section, Laboratory of Immunogenetics, National Institute of Allergy and Infectious Diseases, NIH, Rockville, MD, USA 20852; Molecular Oncology, Department of Medicine, Siteman Cancer Center, Washington University, St. Louis, MO, USA, 63110; Center for Immunology and Inflammatory Diseases, Massachusetts General Hospital and Harvard Medical School, Boston, MA, USA 02129

**Keywords:** Tumor-associated macrophage, antigen presentation, cytotoxic T lymphocyte, cancer

## Abstract

Tumor progression is associated with overstimulation of cytotoxic T lymphocytes (CTLs), resulting in a dysfunctional state of exhaustion. How T cell exhaustion is elicited in the tumor remains poorly understood. Here we show that tumor-associated macrophages (TAMs) present cancer cell antigen and induce CTL exhaustion through a gene expression program dependent on the transcription factor interferon regulatory factor-8 (IRF8). In a transgenic model of murine breast cancer, CTL priming was supported by IRF8-dependent dendritic cells; yet, CTL exhaustion required TAM expression of IRF8, and its ablation suppressed tumor growth. An analysis of the highly immune-infiltrated human renal cell carcinoma tumors revealed abundant TAMs that expressed IRF8 and were enriched for an IRF8 gene expression signature. The IRF8 signature co-segregated with T cell exhaustion markers and was negatively associated with long-term patient survival. Thus, CTL exhaustion is promoted by TAMs via IRF8, and this crosstalk may be disrupted in TAM-targeted therapies.

## Introduction

Malignancy-associated gene mutations can generate neoantigens to activate cancer cell-reactive CD8^+^ cytotoxic T lymphocytes (CTLs). However, CTL effector activities are frequently suppressed in the tumor, with CTLs displaying a dysfunctional state of exhaustion characterized by high expression of the inhibitory receptor programmed cell death 1 (PD-1) (Chen and Mellman, 2017; McLane et al., 2019). Although PD-1 has been successfully targeted for cancer therapy (Baumeister et al., 2016; Sharma and Allison, 2015), how T cell exhaustion is elicited in the tumor remains poorly understood.

Members of the mononuclear phagocyte system (MPS), including monocytes, macrophages and dendritic cells (DCs), function as innate immune effectors and regulators as well as antigen presenting cells (APCs), initiating and modulating adaptive T cell responses. DCs capture antigens in tumor tissues and migrate to lymph nodes to activate antigen-specific T cells. Notably, two classical DC subsets with distinct transcriptional mechanisms of differentiation and APC properties have been identified. DC1s, whose differentiation is dependent on the transcription factor interferon regulatory factor-8 (IRF8), are capable of presenting exogenous antigens to CTLs, while IRF4-dependent DC2s preferentially regulate CD4^+^ T cells (Murphy et al., 2016). In several tumor models, ablation of DC1s impairs the priming of cancer cell-reactive CTLs (Hildner et al., 2008; Roberts et al., 2016; Salmon et al., 2016). However, it is unknown whether cancer drives a distinct DC1 activation state compared to infection, and whether this priming predetermines CTL exhaustion.

An alternative model of CTL exhaustion is centered on the hypothesis that it is acquired following CTL migration to the tumor tissue, triggered by distinct tumor tissue-associated APC activities and/or through APC-independent mechanisms. Notably, monocyte/macrophage lineage of cells constitute a large proportion of leukocytes infiltrated to tumors and have been implicated in suppressing CTL responses in correlation with poor patient outcomes across multiple tumor types (Gentles et al., 2015; Komohara et al., 2014; Zhang et al., 2012). How monocytes and/or macrophages might repress cancer cell-reactive CTLs is not well understood (DeNardo and Ruffell, 2019). They may inhibit CTL proliferation via distinct metabolic activities including depletion of L-arginine and generation of reactive oxygen species (Rodriguez et al., 2004). They also may modulate CTL effector functions through expression of regulatory cytokines such as interleukin-10 (IL-10) and transforming growth factor-β (TGF-β) (Ruffell et al., 2014), or ligands for co-inhibitory receptors such as programmed cell death 1 ligand 1 (PD-L1) (Lin et al., 2018; Tang et al., 2018). In addition, monocytes and/or macrophages may also indirectly suppress CTLs by competing with DCs for cancer cell antigen uptake and dampening the release of immunogenic alarmins from dying cancer cells (Roberts et al., 2017; Zhou et al., 2020). Given that most of these proposed immunosuppressive mechanisms were not assessed for cancer cell antigen-reactive CTLs *in vivo*, it is unclear to what extent these mechanisms are involved in the induction of tumor CTL exhaustion.

A confounding factor in assessing the precise function and mechanism by which monocyte/macrophage lineage of cells control CTL responses in tumor is their apparent heterogeneity. Macrophages constitutively populate developing tissues and respond to inflammatory insults or tissue distress signals by *in situ* proliferation and/or *de novo* differentiation from monocytes (Ginhoux and Jung, 2014; Wynn et al., 2013). In a transgenic model of murine breast cancer, tumor progression triggers monocyte differentiation to a tumor-associated macrophage (TAM) population phenotypically distinct from mammary tissue macrophages (MTMs), which are the predominant macrophage type in healthy mammary glands (Franklin et al., 2014). Although MTMs are also monocyte-derived, TAMs and MTMs are ontologically distinct, with Notch signaling specifically promoting TAM differentiation (Franklin et al., 2014). Notably, TAM accumulation is associated with the induction of CTL exhaustion, and defective TAM differentiation in the absence of Notch signaling attenuates CTL exhaustion and suppresses tumor growth (Franklin et al., 2014). Nonetheless, the potential immunosuppressive mechanisms of TAMs remain to be clarified.

In this study, we expanded our analysis of tumor-induced changes to cells of the MPS in the breast cancer model, and further explored their differentiation and function. We found that TAMs represented the most abundant member of the MPS and expressed high levels of IRF8, which was required for their expansion, maturation, and presentation of cancer cell-associated antigens to CTLs. While CTL priming in tumor-draining lymph nodes was supported by IRF8-dependent DC1s, CTL exhaustion in the tumor was dependent on TAM expression of IRF8, and macrophage-specific ablation of IRF8 suppressed tumor growth. To extend these findings to cancer patients, we focused on the highly immune-infiltrated human renal cell carcinoma (RCC) tumors that were abundantly infiltrated with IRF8-expressing TAMs. Importantly, a macrophage IRF8 gene expression signature was highly associated with T cell exhaustion markers with abundant CTL infiltration and negatively predicted long-term patient survival. These findings suggest that CTL exhaustion is promoted by TAMs via IRF8-dependent APC functions, and such an immunoregulatory module may be targeted for novel cancer immunotherapy.

## Results

### Mononuclear phagocytic cells of the mammary gland are altered in breast cancer

In the MMTV-PyMT (PyMT) model of breast cancer, TAMs have been implicated in promoting tumor progression, with their terminal differentiation defined by expression of the cell adhesion molecule Vcam1 (Franklin et al., 2014; Lin et al., 2001). Ly6C^+^ classical inflammatory monocytes give rise to TAMs via a pathway distinct from that of monocyte-derived MTMs, which are characterized by high expression of the scavenger receptor Mrc1 (Franklin et al., 2014). Conversely, non-classical monocytes are restricted to vasculature and play roles in tumor metastasis, but not in primary tumor growth (Hanna et al., 2015). To assess how cell transformation broadly impacted the MPS, we characterized Ly6C^+^ classical inflammatory monocytes, MTMs, and TAMs as well as Xcr1^+^ DC1s and CD11b^+^ DC2s in the healthy mammary gland and the tumor (Figure S1A). The proportions of TAMs and MTMs among leukocytes were increased and decreased, respectively, in the tumor (Figure 1A), in line with the notion that mutually exclusive tissue niches support MTM and TAM differentiation. DC2 abundance was unchanged, while DC1s were underrepresented among tumor-infiltrating leukocytes (Figure 1A). Notably, compared to DC1s in the healthy mammary gland, a higher fraction of DC1s in the tumor expressed the integrin molecule CD103 (Figure S1B), a marker associated with maturation and trafficking of tissue DC1s to secondary lymphoid organs (Murphy et al., 2016). Indeed, migratory DC1s, but neither resident DC1s nor DC2s in mammary gland- and tumor-draining lymph nodes, expressed CD103 (Figure S1C). These observations reveal that mammary tissue transformation results in accumulation of tumor-resident TAMs as well as maturation of lymph node-homing DC1s.

**Figure 1.**
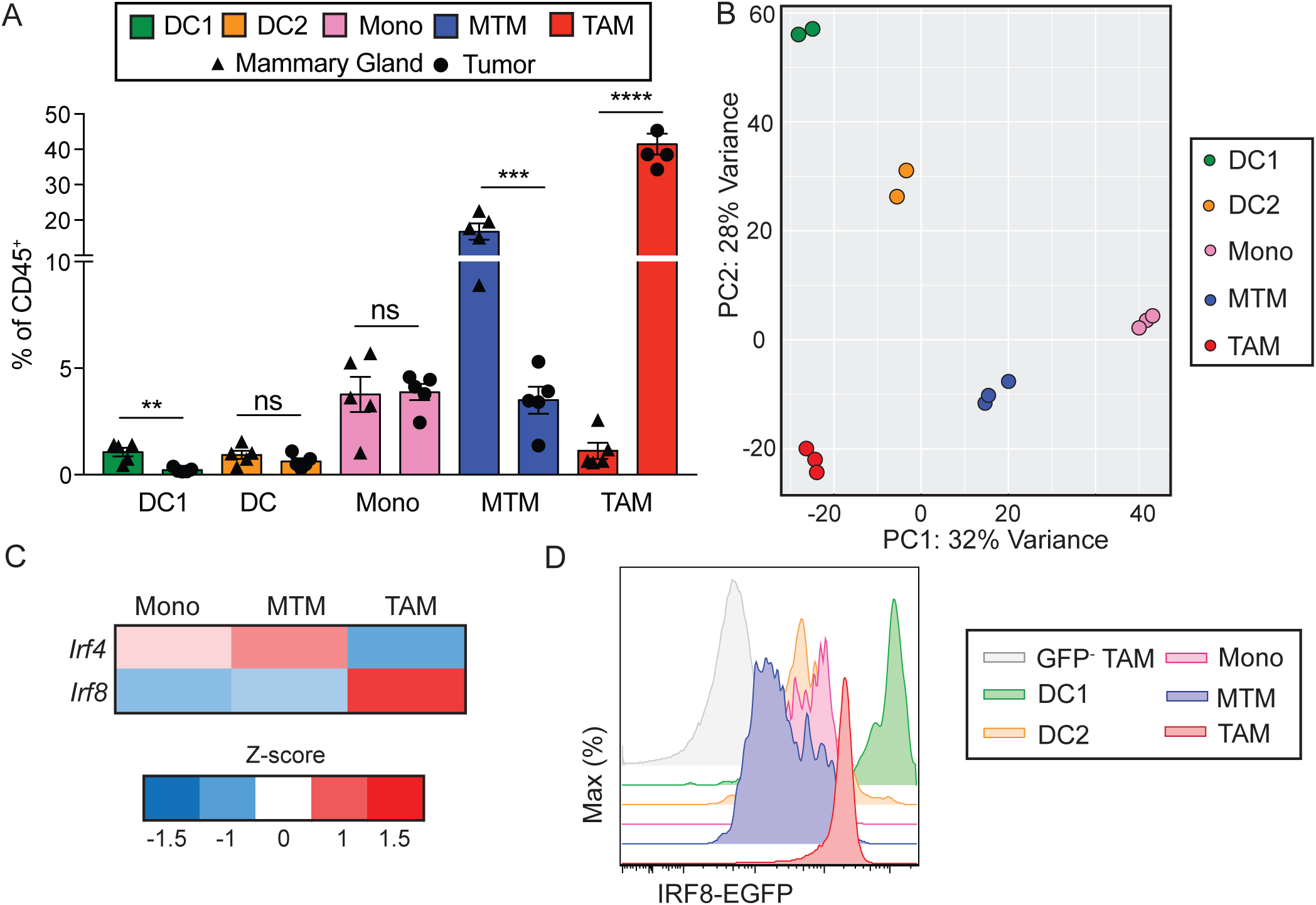
Mononuclear phagocytic cells of the mammary gland are altered in breast cancer. (A) Quantification of DC1s (CD45^+^Lin^-^F4/80^-^Ly6C^-^CD11c^+^MHC-II^+^Xcr1^+^), DC2s (CD45^+^Lin^-^F4/80^-^ Ly6C^-^CD11c^+^MHC-II^+^CD11b^+^), monocytes (CD45^+^Lin^-^F4/80^+^Ly6C^+^CD11b^+^), MTMs (CD45^+^Lin^-^ F4/80^+^Ly6C^-^Mrc1^+^) and TAMs (CD45^+^Lin^-^F4/80^+^Ly6C^-^Mrc1^-^) in mammary glands from virgin mice (triangle) and in tumors of PyMT mice (circle), n=5 per group (Lin = Ly6G, B220, SiglecF, dead cells). (B) Principal component analysis on the transcriptome of DC1, DC2, monocyte, MTM and TAM populations isolated from PyMT tumors, n=2 for DC1, DC2, each circle representing pooled tumors from two mice, n=3 for monocyte, MTM and TAM, each circle representing pooled tumors from one mouse. (C) Expression of two members of the IRF family, IRF4 and IRF8, that are significantly differentially expressed in one or more of the following pairwise comparisons: monocytes versus MTMs, monocytes versus TAMs, and MTMs versus TAMs (base mean expression >100, FDR < 0.05, log_2_ fold change > 0.5. < −0.5, see also Table S1). (D) Flow cytometric analysis of tumor-infiltrating immune cells from IRF8^EGFP^PyMT tumors, which express EGFP as an indicator of IRF8 expression. DC1s (green), DC2s (orange), monocytes (pink), MTMs (blue), TAMs (red) from IRF8^EGFP^PyMT tumor and TAMs from control PyMT tumor (GFP^-^, gray) are shown. Image is representative of five independent experiments. See also Figure S1.

To delineate gene expression programs in the tumor MPS, we performed RNA sequencing (RNAseq) experiments of DC1s, DC2s, monocytes, MTMs, and TAMs isolated from PyMT tumors. Principal component (PC) analyses showed that PC2 separated DC1s and DC2s from monocytes and macrophages, in support of their distinct origin, whereas PC1 segregated monocytes and MTMs from TAMs, DC1s, and DC2s (Figure 1B), revealing shared characteristics among TAMs and DCs. These observations were in line with the finding that TAM terminal differentiation is dependent on RBPJ (Franklin et al., 2014), a transcriptional coregulator of the Notch signaling pathway that also promotes DC maturation (Murphy et al., 2016). To investigate whether TAM differentiation from Ly6C^+^ monocytes was additionally controlled by cell-intrinsic transcriptional programs, we performed differentially expressed gene (DEG) analysis of transcription factors among Ly6C^+^ monocytes, MTMs, and TAMs (Table S1) and focused on the IRF family, which controls various leukocyte fate decisions. Six of the nine IRF family members were differentially expressed among Ly6C^+^ monocytes, MTMs, and TAMs (Table S1), including IRF3, which has previously been associated with TAMs (Biswas et al., 2006). We focused on two members, IRF4 and IRF8, given their opposing expression patterns in MTMs and TAMs (Figure 1C), which resembled antagonism observed in other immune cell lineages, including DC1s and DC2s (Murphy et al., 2016; Singh et al., 2013). IRF8 functions both as a cell lineage-defining transcription factor, as in DC1s and monocyte precursors (Aliberti et al., 2003; Schiavoni et al., 2002; Sichien et al., 2016), and as a mediator of acute responsiveness to extracellular signals, such as to interferon (Langlais et al., 2016). Notably, the interferon target IRF1 was not induced in TAMs relative to monocytes (Table S1), suggesting that IRF8 is upregulated via TAM-intrinsic mechanisms. To validate this pattern of IRF8 expression, we crossed PyMT mice to the *Irf8*^EGFP^ background in which expression of the enhanced green fluorescent protein (EGFP) is under the control of the *Irf8* gene locus. Indeed, TAMs expressed higher EGFP than monocytes or MTMs, although DC1s expressed EGFP at a still higher level (Figure 1D).

### IRF8 deficiency impairs both dendritic cell and macrophage compartments in the tumor

To interrogate the role of IRF8 in the tumor MPS compartments, we crossed a conditional allele of *Irf8* (*Irf8*^fl^) with both CD11c^Cre^ and PyMT transgenic mice, resulting in IRF8 deletion in CD11c^+^ cells, including TAMs, MTMs, and both subsets of DCs (Figure S2A). In line with a specific function for IRF8 in DC1 differentiation (Murphy et al., 2016), there was an approximate 11-fold reduction of tumor DC1s but no impact on DC2 abundance in CD11c^Cre^*Irf8*^fl/fl^PyMT mice (Figure 2A). In addition, there was an almost complete loss of DC1s systemically, in both the resident and migratory DC compartments in tumor-draining lymph nodes (Figure S2B). Notably, a 1.5-fold decrease in total F4/80^+^SiglecF^-^ cells was observed; however, this defect was due neither to Ly6C^+^ monocytes nor to MTMs, as their abundances remained comparable (Figure 2B). Alternatively, there was a 1.9-fold decrease in total TAMs (Figure 2B). Among the TAMs that remained, we observed a 2.2-fold reduction in MHC-II and 1.6-fold reduction in Vcam1 expression levels (Figure 2C), which are markers of TAM terminal differentiation in the PyMT model, revealing that IRF8 is essential for the expansion and maturation of TAMs.

**Figure 2.**
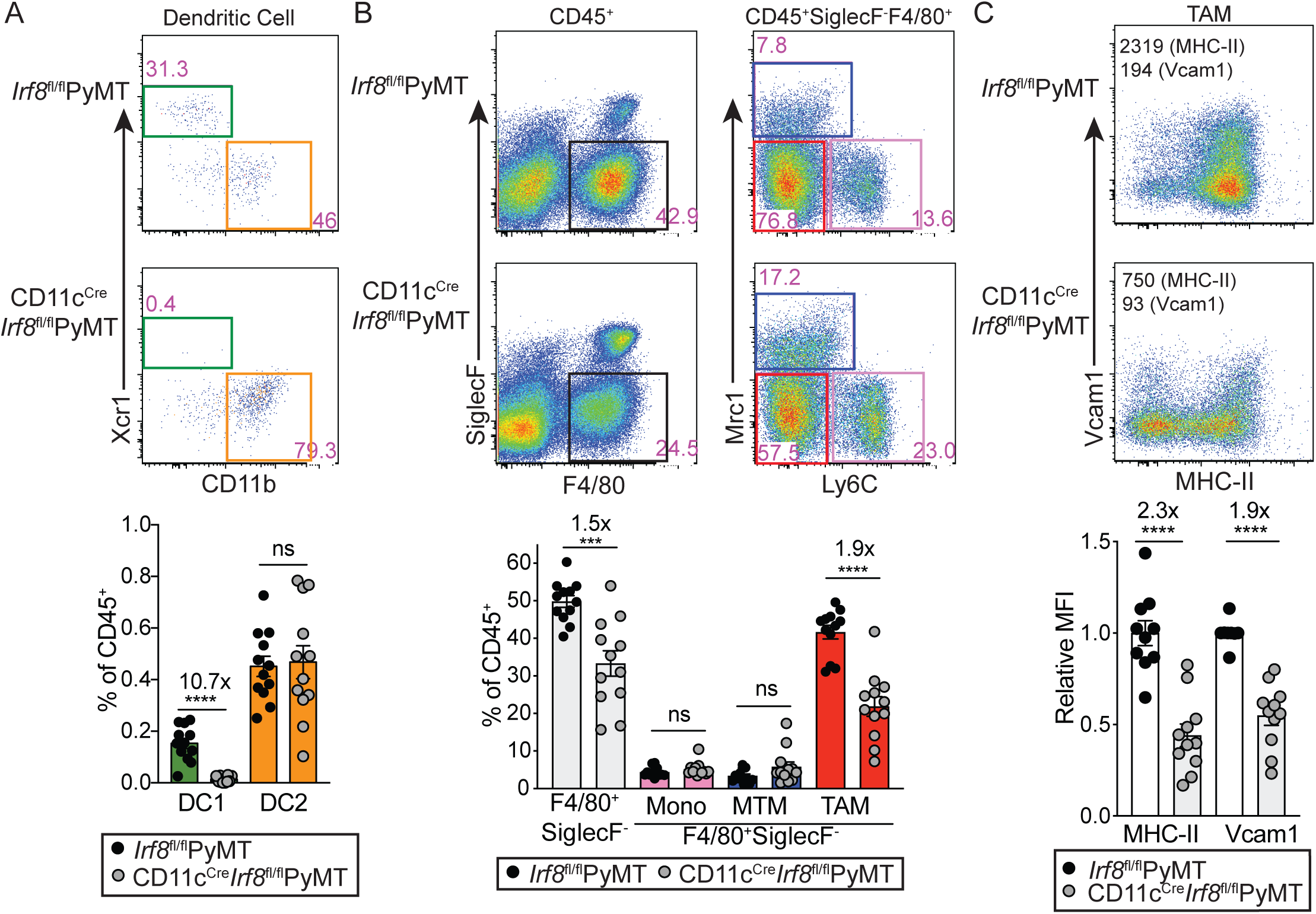
IRF8 deficiency impairs both dendritic cell and macrophage compartments in the tumor. Quantification of tumor-infiltrating myeloid cell populations in *Irf8*^fl/fl^PyMT and CD11c^Cre^*Irf8*^fl/fl^PyMT tumors. Representative flow plots (upper) and quantification (lower) of DCs (A) and monocytes and macrophages (B) are shown. *Irf8*^fl/fl^PyMT (black circles), CD11c^Cre^*Irf8*^fl/fl^PyMT (gray circles), n=12 per group. (C) Representative flow plots and quantification of relative geometric mean fluorescence intensity (MFI) of MHC-II and Vcam1 in CD11c^Cre^*Irf8*^fl/fl^ TAMs (n=11) relative to *Irf8*^fl/fl^PyMT TAMs (n=10), fold calculated within each experiment. Each circle represents pooled tumors from one mouse; comparably sized tumors were selected between *Irf8*^fl/fl^PyMT and CD11c^Cre^*Irf8*^fl/fl^ mice. All data is shown as mean +/− SEM, relative fold change values of means are displayed above significant comparisons (unpaired student’s t test, two-tailed, *** = p<0.001, **** = p<0.0001). See also Figure S2.

Given the disruption in both DC1 and TAM myeloid compartments, we wondered whether the tumor lymphocyte response was impacted. We observed a dramatic loss of CD8^+^ T cells in the tumor, with no difference in abundance of either conventional CD4^+^ cells or Foxp3^+^ regulatory T (Treg) cells (Figure S2C). Although we observed a CTL defect, CD11c^Cre^*Irf8*^fl/fl^PyMT mice showed no difference in tumor growth relative to littermate controls (Figure S2D). This confirms that the CTL response in the PyMT model is ineffective in battling the tumor, and removal of the dysfunctional CD8^+^ T cells does not impact disease outcomes (DeNardo et al., 2009). Given that DC1s are known critical regulators of CTL priming, we were not able to tease apart potential contributions of TAMs to this CTL phenotype, and we aimed to further explore the role IRF8 plays in TAMs.

### IRF8 in TAMs drives a broad antigen presenting cell gene expression program

Given that IRF8 regulates TAM differentiation, we further explored the IRF8-dependent gene expression program by performing RNAseq experiments on TAMs isolated from *Irf8*^fl/fl^PyMT (IRF8 wild-type, WT) and CD11c^Cre^*Irf8*^fl/fl^PyMT (IRF8 knockout, KO) mice. 118 and 92 genes were significantly downregulated and upregulated, respectively, in KO TAMs (Figure 3A and S3A, and Table S2). We applied further FPKM filters to ensure for robust gene expression and generated up- and down-regulated signatures, which included 58 and 53 downregulated and upregulated genes, respectively (Table S2). The IRF8-dependent up-regulated and down-regulated genes were most and least enriched, respectively, in WT TAMs as well as in DC1s and DC2s, while the opposite trend was observed for monocytes and MTMs (Figure 3B and S3B). Additionally, Ingenuity Pathway Analysis revealed that the dendritic cell maturation pathway was enriched in WT TAMs (Figure S3C and Table S3). Notably, when the WT and KO TAM pairs were added to the PC plot, the KO TAMs moved toward MTMs and monocytes but away from WT TAMs and DC1s along PC1 (Figure S3D), supporting the notion that IRF8 drives DC-like properties in TAMs. Importantly, IRF8 deficiency in TAMs did not alter their PC plot position along PC2 (Fig S3D), and the DC-specific transcription factor *Zbtb46* was undetectable in both WT and KO TAMs (data not shown), demonstrating that TAMs acquire DC characteristics in an IRF8-dependent manner while still maintaining their macrophage identity.

**Figure 3.**
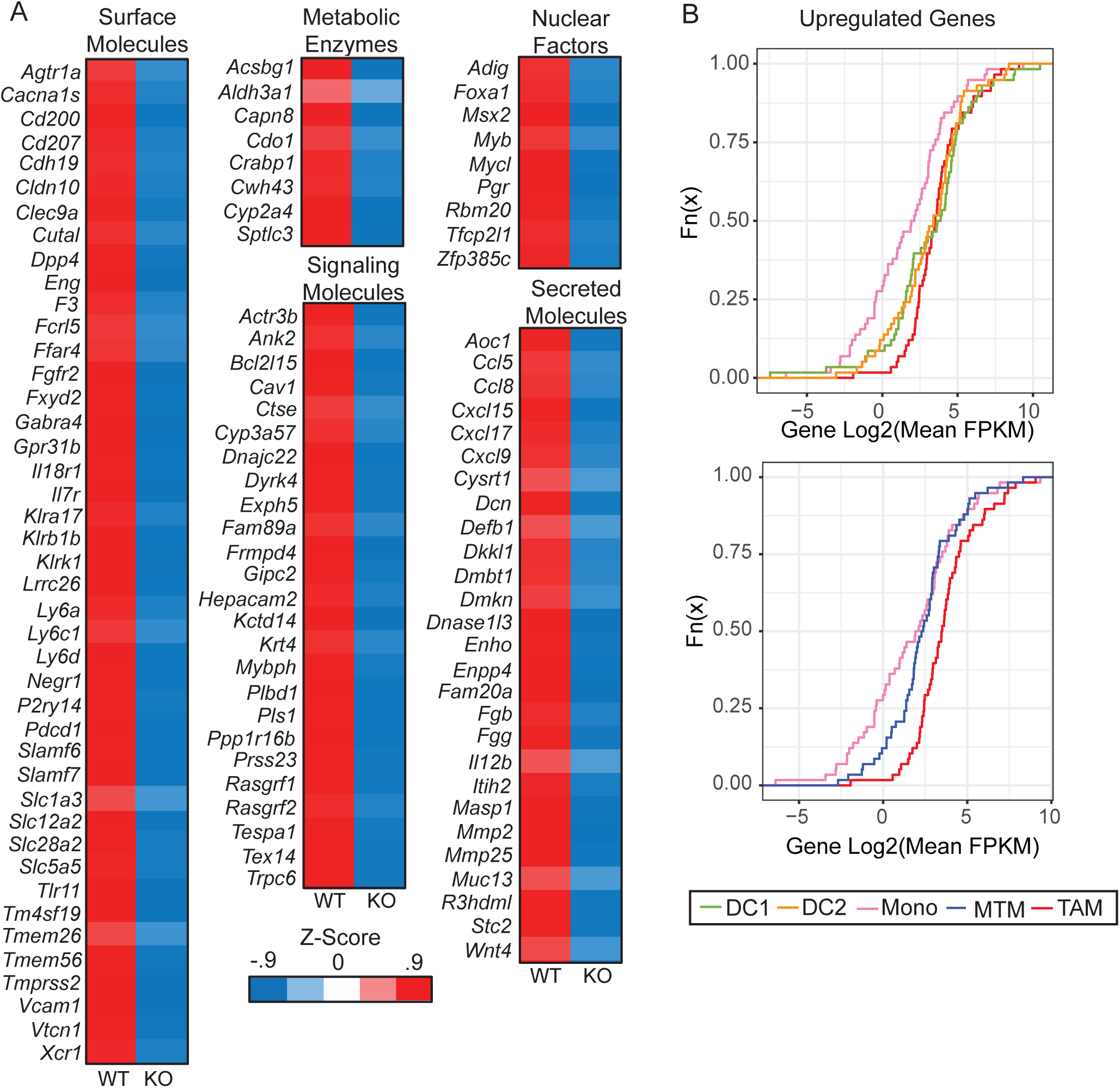
IRF8 in TAMs drives a broad antigen presenting cell gene expression program. (A) Average z-score of genes significantly downregulated in TAMs sorted from tumors of *Irf8*^fl/fl^PyMT (wild-type, WT) and CD11c^Cre^*Irf8*^fl/fl^PyMT (knock-out, KO) mice. RNA was extracted and sequenced. Genes were grouped based on cell localization and function, excluding genes with unknown functions, pseudogenes and noncoding RNAs (*BC035044, BC064078, F630111L10Rik, Gm32014, 2610035D17Rik,* and *A630012P03Rik)*. (B) CDF plot displaying enrichment of IRF8-activated gene signatures in PyMT tumor DC1 (green), DC2 (orange), monocyte (pink), MTM (blue) and TAM (red). See also Figure S3.

Among the downregulated IRF8 target genes in KO TAMs were those encoding cell surface molecules including scavenger and innate immune receptors *Cd207, Clec9a*, *Tlr11* and *Tmprss2* as well as T cell adhesion and regulatory proteins *Cd200, Slamf6, Slamf7*, *Vcam1* and *Vtcn1* (Figure 3A). In addition, genes encoding signaling molecules involved in antigen processing such as *Ctse* as well as an array of targets involved in cytoskeleton and cell mobility control such as *Actr3b, Ank2, Pls1, Rasgrf1* and *Rasgrf2* were downregulated in KO TAMs (Figure 3A). Expression of the transcription factor *Mycl*, a known IRF8 target involved in DC1 priming of T cells (Kc et al., 2014), was also lower in KO TAMs (Figure 3A). Furthermore, in the absence of IRF8, expression of several T cell chemokine genes including *Ccl5, Ccl8, Cxcl9, Cxcl15*, and *Cxcl17* were reduced in TAMs (Figure 3A). Collectively, IRF8 appears to activate a gene expression program that is largely shared with DCs, promoting TAM antigen acquisition and presentation as well as immune cross-talk with T cells.

### TAMs present cancer cell antigen and induce PD-1 expression on CTLs via IRF8

IRF8-dependent DC1s excel in antigen cross-presentation to CD8^+^ CTLs (Murphy et al., 2016). We were intrigued by the finding that IRF8 endows DC-like properties to TAMs and wished to explore whether TAMs could function as APCs and cross-present tumor-associated antigens to CTLs by engineering an *in vivo* tumor antigen presentation system. Gene expression studies revealed that the *S100a8* gene was highly expressed in PyMT cancer cells but not in the untransformed mammary epithelia (data not shown). By crossing S100a8^Cre^ transgenic mice to a PyMT *Rosa26*^LSL-YFP^ background, where the yellow fluorescent protein (YFP) expression cassette is placed downstream of a floxed “stop” sequence (LSL) in the *Rosa26* locus, we confirmed that the Cre recombinase targeted over 80% of cancer cells as well as neutrophils, but not monocytes, MTMs, or TAMs (Figure S4A-B). In order to generate cancer cell-specific antigens, S100a8^Cre^PyMT mice were bred with *LSL-USA* mice, in which a Universal Self Antigen (USA) expression cassette containing multiple model antigen sequences including the SIINFEKL peptide of ovalbumin is driven by the CAG promoter downstream of an LSL sequence (Zhang et al., 2019). Before tumors grew, S100a8^Cre^*LSL-USA*PyMT mice were lethally irradiated and reconstituted with congenically mismatched bone marrow cells from wild-type (WT) and CD11c^Cre^*Irf8*^fl/fl^ (KO) mice mixed at a 1:1 ratio (Figure 4A). The chimeric mice were aged to allow bone marrow cells to graft and tumors to grow, at which point endogenous neutrophils were replaced by donor bone marrow (data not shown), leading to tumor-restricted antigens. The capabilities of these WT and KO TAMs to cross-present cancer cell-specific antigens picked up *in vivo* could be compared via coculture with SIINFEKL-reactive CD8^+^ OT-1 T cells *in vitro* (Figure 4A).

**Figure 4.**
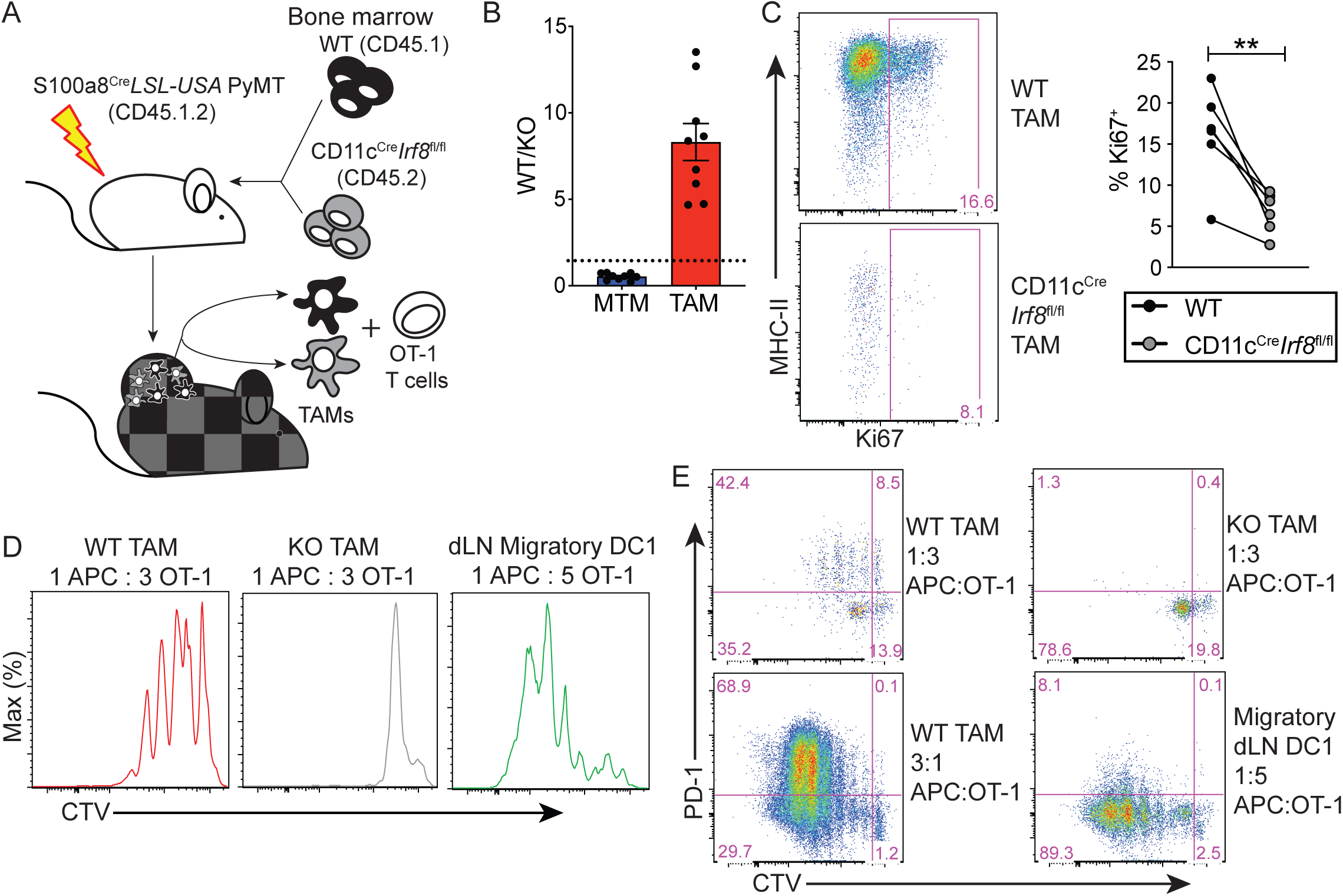
TAMs present cancer cell antigen and induce PD-1 expression on CTLs via IRF8. (A) Schematic showings mixed bone marrow chimera experimental setup, with recipient mice (S100a8^Cre^*LSL-USA*PyMT, CD45.1.2) and bone marrow transfer (1:1) of C57BL/6 CD45.1 (WT, black) and CD11c^Cre^*Irf8*^fl/fl^ CD45.2 (KO, gray) bone marrow. TAMs were sorted from tumors of chimeric mice and cocultured with naïve OT-1 CD8^+^ T cells. (B) Reconstitution of TAMs in tumors of chimeric mice. After aging chimeric mice described in (A), and upon sizeable tumor burden, tumor-infiltrating immune cells were quantified. Ratios of WT to KO MTMs and TAMs were calculated, each dot representing pooled tumors from one mouse (n=9). (C) Representative flow cytometric gating and quantification of Ki67 in WT and KO TAMs from chimeric mice (n=6), paired student’s t-test, two-tailed, ** = p<0.01. (D) Ability of WT and KO TAMs from tumor and WT migratory DC1s from tumor-draining lymph nodes (dLN) to trigger proliferation of OT-1 CD8 T cells. WT and KO TAMs were sorted from tumors and WT migratory DC1 (CD45^+^MHCII^high^CD11c^+^Xcr1^+^) were sorted from dLN of chimeric mice, cocultured with CTV-labeled naïve OT-1 CD8^+^ T cells at indicated ratios in T cell media at indicated cell ratios. T cells were analyzed by flow cytometry 72 hours later. (E) PD-1 expression from coculture experiments shown in D as well as a higher TAM concentration condition. Data representative of 3 independent experiments. See also Figure S4.

Upon initial characterization of the macrophage compartments, KO bone marrow cells contributed to TAMs approximately 8-fold less than WT bone marrow cells, while differentiation into MTMs by KO cells was enhanced (Figure 4B and S4C). This was coupled with a loss of Ki67 expression in KO TAMs (Figure 4C). These observations corroborate the conclusion that IRF8 specifically promotes the differentiation, expansion, and/or fitness of TAMs (Figure 2). WT and KO TAMs were further isolated from tumors of chimeric mice, cocultured with OT-1 T cells labeled with the cell proliferation dye CellTrace Violet (CTV) for 3 days. WT, but not KO, TAMs induced substantial OT-1 T cell proliferation, although the T cell stimulatory activity of TAMs was less potent than that of WT migratory DC1s isolated from the tumor-draining lymph nodes (Figure 4D and S4D). Notably, induction of PD-1 expression was observed in OT-1 T cells cocultured with WT TAMs at multiple concentrations, leading to varying degrees of proliferation, whereas those cocultured with KO TAMs or DC1s were much less efficient in gaining PD-1 expression (Figure 4E). These findings reveal that tumor-resident TAMs can cross-present cancer cell-associated antigens to CTLs, and induce T cell exhaustion in an IRF8-dependent manner.

### Macrophage-specific deletion of IRF8 attenuates CTL exhaustion in the tumor

We considered the possibility that a requirement for DC1-dependent priming of CD8^+^ T cells could mask IRF8-dependent CTL regulatory functions of TAMs in the tumor of CD11c^Cre^*Irf8*^fl/fl^PyMT mice. To this end, we generated MafB^iCre^*Irf8*^fl/fl^PyMT mice, resulting in IRF8 deletion in macrophages and monocytes, but not in DCs (Wu et al., 2016) (Figure S5A). These mice displayed intact DC compartments both in the tumor-draining lymph nodes and in the tumor (Figure 5A and S5B-C). In contrast, TAMs from MafB^iCre^*Irf8*^fl/fl^PyMT mice displayed the same defect as that of CD11c^Cre^*Irf8*^fl/fl^PyMT mice, with a decrease in abundance, maturation, and proliferation, as determined by expression of MHC-II, Vcam1, and Ki67 (Figure 5B-C, S5D and data not shown). In contrast to CD11c^Cre^*Irf8*^fl/fl^PyMT mice, the abundance of CD8^+^ T cells in MafB^iCre^*Irf8*^fl/fl^PyMT mice was comparable to that of *Irf8*^fl/fl^PyMT mice (Figure 5D and S5E-F). These observations suggest that DC1s are critical regulators of the CTL response in the tumor, presumably through priming in the tumor-draining lymph node (Hildner et al., 2008; Roberts et al., 2016; Salmon et al., 2016), whereas TAM IRF8 is negligible for modulating the magnitude of this responses.

**Figure 5.**
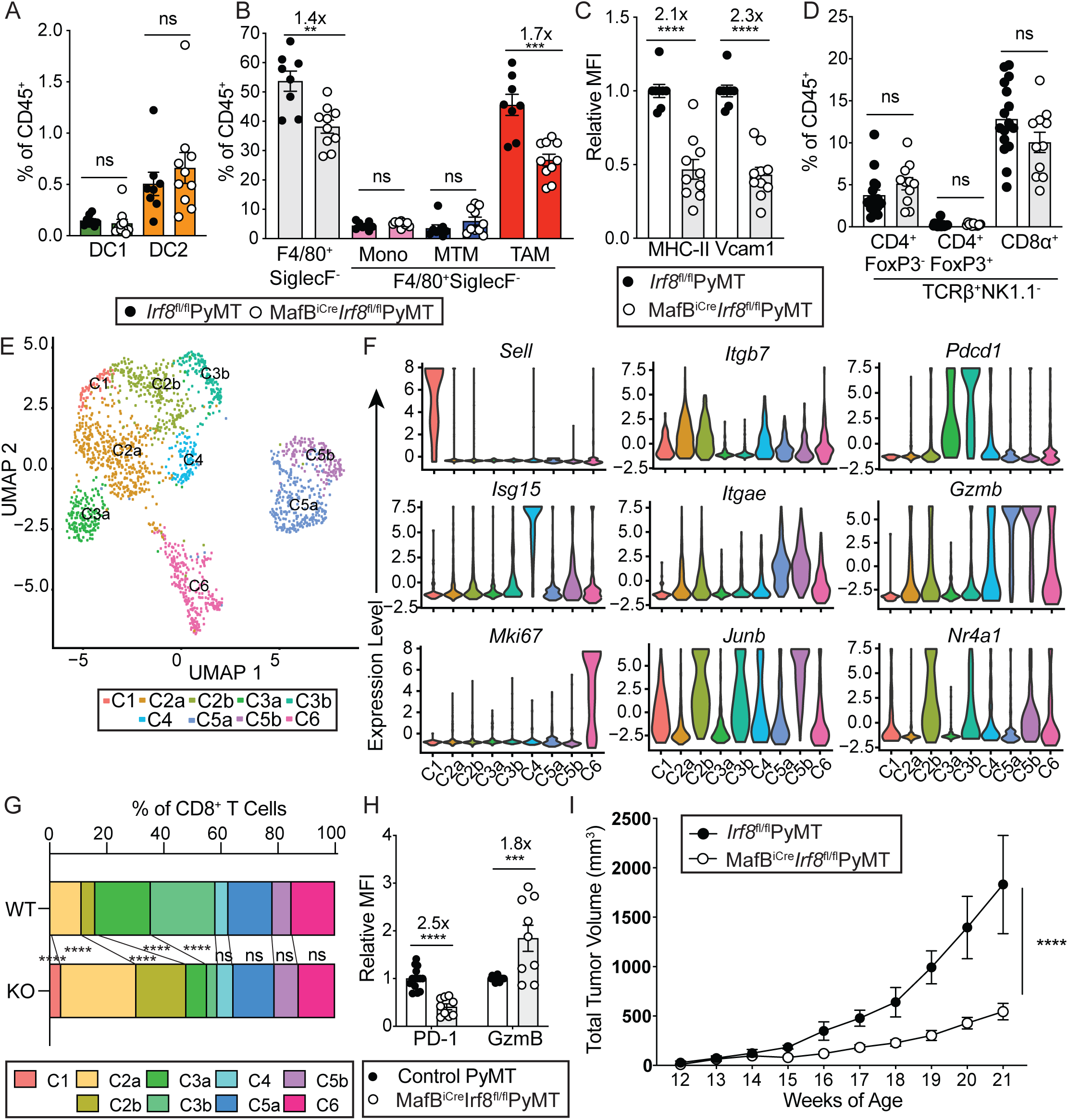
Macrophage-specific deletion of IRF8 attenuates CTL exhaustion in the tumor. (A) Quantification of tumor-infiltrating dendritic cells in *Irf8*^fl/fl^PyMT (WT, black circles, n=8) and MafB^iCre^*Irf8*^fl/fl^PyMT (KO, white circles, n=10) mice. DC1 (green), and DC2 (orange) are shown. (B) Quantification of tumor-infiltrating monocyte/macrophage lineage of cells in WT (black circles, n=8) and KO (white circles, n=10) mice. Total F4/80^+^SiglecF^-^ cells (gray), monocytes (pink), MTMs (blue), and TAMs (red) are shown. (C) Quantification of relative geometric mean fluorescence intensity (MFI) of MHC-II and Vcam1 relative to WT TAMs, MFI indicated on representative plot (WT n=8, KO n=10). (D) Quantification of tumor-infiltrating conventional and regulatory CD4^+^ and conventional CD8^+^ T cells in *Irf8*^fl/fl^PyMT (n=16) and MafB^iCre^*Irf8*^fl/fl^PyMT (n=11) mouse tumors. (E) UMAP projection of single-cell RNA sequencing (scRNAseq) clusters of TCRβ^+^CD8α^+^ cells sorted from multiple tumors of one *Irf8*^fl/fl^PyMT and one MafB^iCre^*Irf8*^fl/fl^PyMT mouse. (F) Violin plots showing normalized expression of select genes across all 9 clusters from scRNAseq dataset. (G) Abundance of each cluster as a proportion among total cells within WT or KO tumors. (H) Relative MFI of PD1 and Granzyme B (GzmB) of TCR*b*^+^NK1.1^-^CD8*a*^+^ T cells from tumors of WT and KO mice. (I) Tumor measurements from MafB^iCre^*Irf8*^fl/fl^PyMT and littermate and cagemate *Irf8*^fl/fl^PyMT controls (n=7 each). All data are shown as mean +/− SEM, fold change of means are displayed above significant comparisons. A, B, C, D, H: each circle represents pooled tumors from one mouse; comparably sized tumors were selected between WT and KO mice, unpaired student’s t-test, two-tailed, ** = p<0.01, *** = p<0.001, **** = p<0.0001. H: Fisher’s Exact Test, **** = p<0.0001 I: 2-way ANOVA **** = p<0.0001. See also Figure S5 and S6.

While we did not observe differences in overall CTL abundance, we wanted to investigate whether TAM IRF8 influences CTL phenotypes. To this end, we performed single-cell RNA sequencing (scRNAseq) experiments of tumor-infiltrating CD8^+^ T cells from *Irf8*^fl/fl^PyMT (WT) and MafB^iCre^*Irf8*^fl/fl^PyMT (KO) mice, which revealed 9 clusters and 6 broad phenotypes (Figure 5E and S6). Naïve-like T cells, exemplified by expression of *Sell*, were represented by C1. Early-activated and effector CD8^+^ T cells, which expressed high levels of *Itgb7* and intermediate levels of *Gzmb* and *Itgae*, were found in C2a and CD2b, while exhausted cells, identified by high *Pdcd1* expression, were found in C3a and C3b. Other clusters included cells with an interferon signature, expressing *Isg15*, in C4, innate-like T cells (ILTCs), identified previously (Dadi et al., 2016), in C5a and C5b, and proliferating cells expressing *Mki67* in C6 (Figure 5F). Multiple phenotypic categories were broken into two clusters, differentiated by expression of immediate-early genes downstream of T cell receptor (TCR) signaling including *Junb* and *Nr4a1* (C2b, C3b, C5b) (Figure 5E-F and Table S4). Thus, a subset of effector, exhausted, and ILTCs acutely see antigen in the tumor microenvironment, which may come from DCs, TAMs, or cancer cells themselves.

While CD8^+^ T cells from WT and KO mice were represented by the same clusters, the relative abundance of specific clusters were altered in KO mice, with enrichment for naïve-like and early-activated/effector T cells (C1, C2a, C2b) and diminished exhausted clusters (C3a, C3b) (Figure 5G). In accordance with the altered abundances, tumor-infiltrating CD8^+^ T cells showed increased granzyme B and decreased PD-1 protein expression via flow cytometric analysis, both on a population level and by geometric mean fluorescence intensity (Figure 5H and S5G). Notably, this altered CTL compartment was associated with tumor protection, as MafB^iCre^*Irf8*^fl/fl^PyMT mice exhibited slower tumor growth than littermate controls (Figure 5I). These observations suggest that in the PyMT model, while DC1s are required for initial priming of tumor-specific CD8^+^ T cells, re-encountering antigen on IRF8-expressing TAMs in the tumor promotes T cell exhaustion, thus attenuating CD8^+^ T cell cancer immunosurveillance.

### TAMs from human clear cell RCC patients express IRF8 and IRF8-activated genes

We wished to determine whether a similar TAM-dependent immune regulatory module occurred in human solid tumors. We chose to investigate clear cell renal cell carcinoma (ccRCC), as it is a highly immunogenic tumor type with robust T cell and macrophage responses (Chevrier et al., 2017). Furthermore, with surgery being the first line therapy, naturally evolved tumor-elicited immune responses can be observed in drug treatment-naïve cancer patients. Surgically removed fresh tumor tissues underwent flow cytometric analysis to investigate MPS tumor infiltration, including DC1s, DC2s, classical monocytes, and TAMs, defined as HLADR^+^CD14^-^CD141^+^, HLADR^+^CD14^-^CD1c^+^, CD14^+^HLADR^low^, and CD14^+^HLADR^high^ leukocytes, respectively (Figure 6A). As in mice, DC1s and DC2s were very rare, making up less than one percent of the immune infiltrate, whereas TAMs were 31-fold more abundant than DC1s and 19-fold more abundant than DC2s (Figure 6A). The IRF8 expression pattern across MPS compartments was similar to that observed in mice, with DC1s expressing the highest level and TAMs expressing the second highest level (Figure 1D and 6B). Additionally, in most patient samples, we observed higher IRF8 expression in TAMs than in monocytes, again matching what we observed in our mouse tumor model (Figure 1D and 6C).

**Figure 6.**
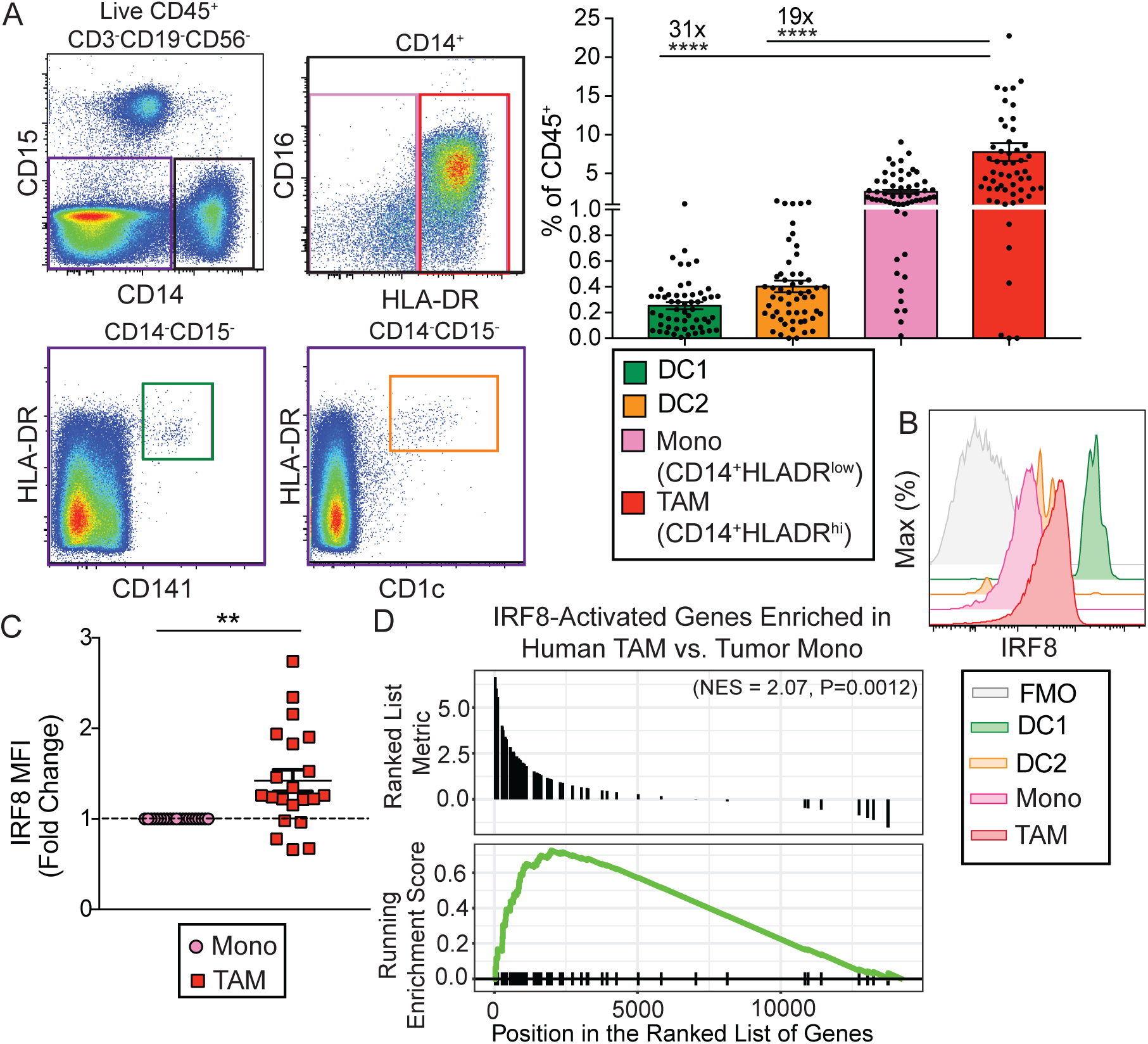
TAMs from human clear cell RCC patients express IRF8 and IRF8-activated genes. (A) Representative flow plots and quantification of tumor-infiltrating DC1s (green, CD45^+^Lin^-^CD14^-^ HLA-DR^+^CD141^+^), DC2s (orange, CD45^+^Lin^-^CD14^-^HLA-DR^+^CD1c^+^), classical monocytes (pink, CD45^+^Lin^-^CD14^+^HLA-DR^low^), and TAMs (red, CD45^+^Lin^-^CD14^+^HLA-DR^high^) in human clear cell RCC tumor samples as determined by flow cytometric analysis (Lin = CD15, CD19, CD3, CD56, dead cells). Fold differences in abundance between TAMs and DC1s, TAMs and DC2s are shown. Each circle represents one patient tumor sample, data shown as mean +/− SEM, n=56 (one-way ANOVA with multiple comparisons, **** = p<0.0001). (B) Representative plot showing expression of IRF8 in tumor-infiltrating DC1s (green), DC2s (orange), monocytes (pink), TAMs (red), and fluorescence minus one (FMO) TAM sample (gray), as determined by flow cytometric analysis. (C) IRF8 MFI fold change in TAMs relative to tumor-infiltrating monocytes. IRF8 MFI is relative to monocyte MFI (n=21), data shown as mean +/− SEM (paired student’s t test, two-tailed, ** = p<0.01). (D) Enrichment of human orthologs of IRF8-activated mouse TAM genes among human TAM-specific genes relative to human monocytes. Tumor monocytes (n=5) and TAMs (n=6) were sorted from human RCC samples, underwent RNA sequencing and differential gene expression analysis. The significantly upregulated IRF8-activated genes identified in mouse WT TAMs were converted to their human orthologs, and the enrichment of these IRF8-activated genes was tested among human TAM-specific genes.

To define the genetic program that may control TAM differentiation from monocytes, we performed RNAseq on purified monocyte and TAM populations from RCC patient tumors. Significant enrichment of the human orthologs of mouse IRF8-activated target genes was observed in human TAMs over monocytes (Figure 3A and 6D), suggesting that IRF8 induces a similar gene expression program in human TAMs. Thus, we were able to identify TAMs in human solid tumors that expressed both IRF8 and an IRF8-dependent gene program that mirrored what we observed in mouse.

### A TAM IRF8 gene signature predicts CTL exhaustion and RCC patient survival

We aimed to generate a human IRF8-TAM gene signature using the genomics data we had generated from both mouse and human TAMs. After converting the mouse IRF8-activated genes to their human orthologues (Figure 6D), we applied further FPKM filters to ensure robust expression in the human TAM RNAseq dataset. The resulting IRF8-TAM signature consisted of 18 genes (Figure 7A). To explore the specificity of this signature, we looked for its enrichment in RNAseq datasets generated from a number of purified myeloid and lymphoid cell populations from RCC tumors, demonstrating that the signature was most enriched in TAMs, DC1s, and DC2s (Figure 7B). This pattern of enrichment suggests this signature is shared among multiple mature APC compartments in human RCC tumors, similar to what we observed in mice (Figure 3B).

**Figure 7.**
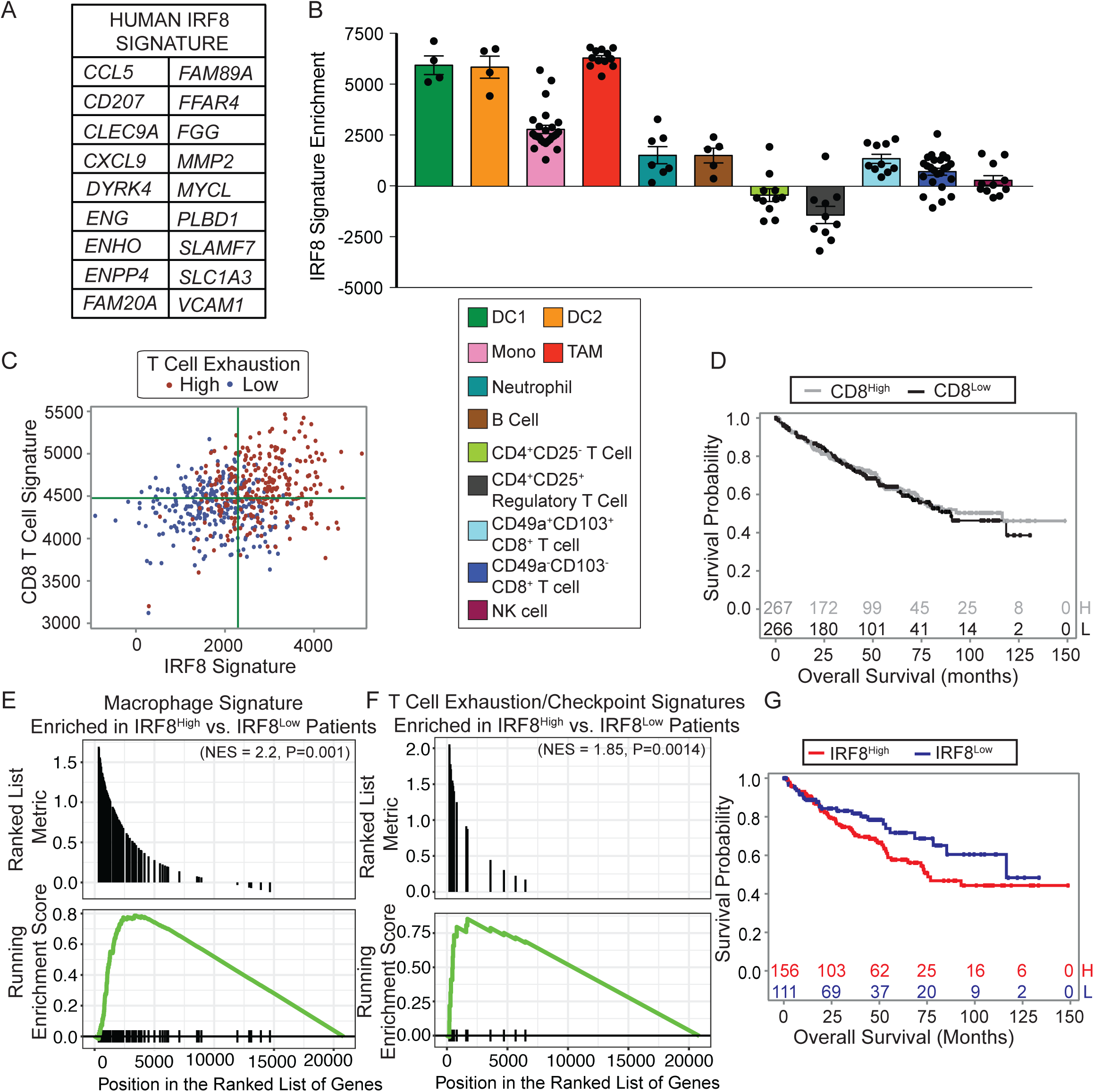
A TAM IRF8 gene signature predicts CTL exhaustion and RCC patient survival. (A) Human IRF8 signature was generated by using additional FPKM filters (FPKM mean >1, FPKM variance > 3) on human orthologs of IRF8-activated mouse TAM genes. (B) Enrichment of Human IRF8 Signature across various immune populations sorted from tumor and peripheral blood of RCC patients, including DC1s, DC2s, monocytes, TAMs, neutrophils (CD15^+^CD16^+^), B cells (CD19^+^), conventional CD4^+^ T cells (CD3^+^CD4^+^CD25^-^), Treg cells (CD3^+^CD4^+^CD25^+^), CD49a^+^CD103^+^ CD8^+^ T cells (CD3^+^CD8*a*^+^CD49a^+^CD103^+^), CD49a^-^CD103^-^ CD8^+^ T cells (CD3^+^CD8*a*^+^CD49a^-^CD103^-^), and NK cells (CD3^-^CD56^dim^CD16^+^) (within a cell type: n=4-21, each sample from a different patient). (C) Individual RCC patient enrichment values for IRF8 signature (x-axis) and CD8 T Cell signature (y-axis), green line indicates median values. CD8 T Cell exhaustion signature (red = “high”, top half enrichment, blue = “low”, bottom half enrichment) is also shown. Each dot represents one patient. (D) Survival probability for RCC patients stratified based CD8 T cell signature enrichment in bulk tumor RNAseq (top quadrants versus bottom quadrants in C) (CD8^High^ n=267 and CD8^Low^ n=266). (E) Mac_CSF1 gene set enrichment analysis in IRF8^high^ patients. Bulk tumor RNAseq from a cohort of RCC patients was analyzed (n=533). Patients were ranked based on enrichment for the human IRF8 signature, split into IRF8^High^ and IRF8^Low^, and enrichment of the macrophage signature was tested. (F) Same as in (E), except enrichment of pooled CD8 T cell checkpoint/exhaustion gene signatures was tested. (G) Survival probability for IRF8^High^ and IRF8^Low^ patient cohorts within CD8^high^ patients (top left and top right quadrants in C) (CD8^high^ total: n=267, IRF8^High^: n=156, IRF8^Low^: n=111, p=0.08).

To investigate whether the IRF8 signature was associated with phenotypically distinct CTL responses in tumor, we used the bulk ccRCC patient RNAseq datasets from the TCGA database. We subsetted patients based on enrichment for both a CD8^+^ T cell signature (Bindea et al., 2013) and the IRF8 signature (Figure 7C). The degree of tumor CD8^+^ T cell infiltration did not impact disease outcomes (Figure 7D). The bulk RNAseq from tumors among the IRF8^high^ group was enriched for multiple macrophage signatures (Beck et al., 2009; Bindea et al., 2013) (Figure 7E and data not shown), confirming that TAMs manifest an IRF8-dependent gene expression program and are the major APC subset in the tumor (Figure 6B). In addition, the IRF8^high^ group had significant enrichment for T cell checkpoint and exhaustion signatures (Chung et al., 2017) (Figure 7C and F). Notably, among CD8^+^ T cell-enriched patients, IRF8^high^ patients had worse long-term survival outcomes than IRF8^low^ patients (Figure 7C and G). These observations suggest that in solid tumors with robust CD8^+^ T cell infiltration, the IRF8-dependent TAM gene program promotes CD8^+^ T cell exhaustion in both mouse and human (Figure S7).

## Discussion

Tumor progression is associated with overstimulation of cancer cell-reactive CTLs, resulting in a dysfunctional state of exhaustion. Using murine models of myeloid cell-specific ablation of IRF8, our studies revealed a critical role for TAMs in induction of CTL exhaustion via their abiliy to present cancer cell-associated antigens to CD8^+^ T cells. While APC-independent mechanisms of suppression in the tumor microenvironment have been proposed, we did not observe IRF8-dependent expression of genes implicated in such pathways in TAMs. Given that there is no difference in tumor growth in CD11c^Cre^*Irf8*^fl/fl^PyMT mice, where DC1-primed T cells are absent, any potential CTL-independent pro-tumor TAM activities are not dependent on IRF8 and/or do not contribute in this model. Our findings confirm that IRF8-dependent TAM APC activity is a key regulator of the CTL response

T cells can encounter cancer cell-associated antigens on a variety of cell types. These APC compartments have different localizations, migratory abilities and functions. While DC1s are very rare in the tumor itself, as observed in both mouse and human solid tumors (Broz et al., 2014), they are able to migrate to the draining lymph nodes. It is this migratory ability that allows them to carry out a critical role in priming CTL responses (Roberts et al., 2016; Salmon et al., 2016), and their APC function in the tumor appears negligible (Diao et al., 2018). Conversely, TAMs are highly abundant in solid tumors and are tissue-resident, which may render them the more relevant APC subset to modulate CTL responses in the tumor. While CTLs may encounter directly-presented antigen on cancer cells themselves, our results suggest this CTL-cancer cell interaction is not sufficient to drive exhaustion. This is in contrast to ILTCs (Dadi et al., 2016), which require neither DC1s nor TAMs for their activity in the tumor as we observed, and instead can directly respond to cancer cells (Chou C, et al., unpublished observations). Thus, distinct APC compartments control different types of T cell responses in the tumor.

Our study also confirms differences between DC1 and TAM antigen presenting capabilities to CD8^+^ T cells. DC1s exhibit robust cross-presenting capability and a potent ability to activate these cells, while TAMs are able to present exogenous cancer cell-associated antigen with lower potency in association with their ability to induce CTL exhaustion. How TAMs acquire and/or process exogenous antigens to present on class I major histocompatibility complex (MHCI) remains unclear. TAMs may cross-present cancer cell antigens following engulfment of cancer cell constituents. When assessing the IRF8-dependent targets in TAMs, we do not observe the canonical cross-presentation genes that have been implicated in DC1s. The process of cross-presentation itself is not fully understood, and it remains possible TAMs are partaking in a non-canonical cross-presentation process that is distinct from DC1s (Blander, 2018). In addition, it is possible that TAMs employ “cross-dressing” by capturing preformed peptide-MHCI complexes from cancer cells (Campana et al., 2015). The relative contribution of cross-presentation and cross-dressing in TAM presentation of cancer cell antigens warrants further investigation and can be differentiated by animal models with selective ablation of MHCI antigen presentation genes in TAMs or cancer cells.

The APC function of TAMs requires a robust DC response, as priming of CTLs is required before T cells can traffic to the tumor and reencounter antigen in the context of a TAM. Although our murine cancer model exhibits reproducible and a homogeneous immune response across mice, human patients demonstrate considerable heterogeneity in their tumor immune responses. We focused on ccRCC patients who had a robust DC response, as indicated by CD8^+^ T cell infiltration in the tumor, to then uncover the association of TAM-expressed IRF8 with increased T cell exhaustion and poor disease outcomes. Whether patients have a DC1-dependent priming of cancer cell-reactive CD8^+^ T cells and if that is coupled with a TAM response in the tumor may be important drivers of how CTLs combat cancer.

Monocytes and macrophages are highly adaptable to the tissue environment, and macrophages can be heterogeneous across different organs and tumor types as well as between individuals. The finding that IRF8 and IRF4 are selectively enriched in TAMs and MTMs raises the possibility that analogous to DC1s and DC2s, a division of labor exists among macrophage populations. In some murine cancer models, tissue-resident macrophages derived from embryonic progenitors have been implicated in tumor regulation (Loyher et al., 2018; Zhu et al., 2017). Resident macrophages in multiple organs, including microglia of the brain and macrophages of the kidney and liver express IRF8, and IRF8 plays a critical function in macrophage maturation and survival (Hagemeyer et al., 2016). It will be interesting to determine whether embryonically-derived macrophages also engage the IRF8-dependent antigen presentation program to induce CTL exhaustion in cancer or other conditions of chronic antigen stimulation, as there is evidence Kupffer cells in the liver can cross-present antigen in acute infection (Ebrahimkhani et al., 2011). Much of the work defining embryonically-derived resident macrophages across tissues has been conducted in mice, and it remains unclear how to differentiate these subsets from monocyte-derived macrophages in human. While our work uncovers IRF8 as a commonly expressed marker in human ccRCC TAMs, it is not observed in all patients and there may be further heterogeneity among the TAM phenotypes across human tumors. Further investigation is needed to identify additional markers and clarify the heterogeneity in both origin and function of human TAMs.

In summary, our work has uncovered a key role for IRF8 in driving TAM identity and T cell exhaustion in both mouse and human tumors. Such an immune regulatory function of macrophages may have evolved to prevent collateral tissue damage but is maladapted in the tumor. Current immunotherapies targeting TAM responses under development may work in part through disrupting this crosstalk.

## Acknowledgements

We acknowledge the use of the Integrated Genomics Operation Core, funded by the NCI Cancer Center Support Grant (CCSG, P30 CA08748), Cycle for Survival, and the Marie-Josée and Henry R. Kravis Center for Molecular Oncology. This work was supported by the National Institutes of Health (F31 CA210332 to B.G.N. and R01 CA198280 to M.O.L.), the Howard Hughes Medical Institute (Faculty Scholar Award to M.O.L.), Alan and Sandra Gerry Metastasis and Tumor Ecosystems Center (M.O.L.), the Ludwig Center (A.A.H), the Parker Institute (A.A.H.) and the Weiss Family Foundation (A.A.H).

## Author Contributions

B.G.N. and M.O.L. were involved in all aspects of this study, including planning and performing experiments, analysis and interpretation of data and writing the manuscript. F.K. processed and analyzed all sequencing data and wrote the manuscript. A.A.H. lead and oversaw all human data collection and analyses. M.L., K.C., M.D., E.R.K., R.M.S., T.A.P., C.K., A.S., L.V., and R.A.F. assisted with mouse colony management and performed experiments. X.W., H.C.M.3^rd^, K.M.M., and J.J.M. provided key mouse lines utilized in the studies. J.J.H., T.A.C., A.A.H. and F.K. were involved in initiating, overseeing and planning collaborations regarding human tumor sample collection and analyses.

## STAR Methods

### Lead Contact and Materials Availability

Further information and requests for resources and reagents should be directed to and will be fulfilled by the Lead Contact, Dr. Ming Li. This study did not generate new unique reagents. The raw sequencing data that support the findings of this study are available from the lead contact upon reasonable request.

### Experimental Model and Subject Details

#### Animal experimental models

CD45.1^+^, *Irf8*^fl/fl^, OT-1, and S100a8^Cre^ were purchased from Jackson Laboratory. IRF8^EGFP^ and MafB^iCre^ mice were generated as previously reported (Wang et al., 2014; Wu et al., 2016). *LSL-USA* mice were generated in the laboratory of Dr. James Moon (Zhang et al., 2019). MMTV-PyMT, CD11c^Cre^ and *Rosa26*^LSL-YFP^ were in the lab as previously reported (Franklin et al., 2014). All mice used in these studies were on the C57Bl/6 background and were female given the focus on the mammary gland and PyMT breast cancer. All mice were maintained under specific pathogen-free conditions, and animal experimentation was conducted in accordance with procedures approved by the Institutional Animal Care and Use Committee of MSKCC.

#### Human experimental models

Informed consent was obtained from ccRCC patients prior to surgery utilizing MSKCC IRB protocol 12-237. Tissue was processed on the same day as surgery. The cohort of ccRCC patients from which tissues were analyzed were 91.6% male, mean age 58.3, median age 58.

### Method Details

#### Tumor measurements

Beginning at 12 weeks of age, tumors were measured weekly with a caliper. Tumor volume was calculated using the equation [(LxW^2^) x (π/6)], in which L denotes length and W denotes width. Individual tumor volumes were added together to calculate total tumor burden, with an endpoint when total volume reached 3,000 mm^3^ or one tumor reached 2,000 mm^3^. Researchers were blinded to genotypes of mice during measurements.

#### Immune cell isolation from tissues

Immune cells were isolated from mouse mammary gland, mouse tumors and human tumors as described previously (Dadi et al., 2016). Briefly, tissues were prepared by mechanical disruption with a razor blade followed by incubation in 280 U/mL Collagenase Type 3 and 4 μg/mL DNase I in HBSS at 37°C for 1 h with periodic vortex. Digested tissues were passed through 70 μm filters and pelleted. Cells were resuspended in 44% Percoll and layered above 66% Percoll. Sample was centrifuged at 1,900 g for 30 min without brake. Cells at interface were collected, stained with antibodies and analyzed by flow cytometry or used for FACS. To isolate cells from tumor-draining lymph nodes, tissues were mechanically disrupted with a razor blade, then incubated in 300 U/mL Collagenase Type 3 in RPMI supplemented with 10% FBS for 35 min at 37°C. EDTA was added for final concentration of 0.01 M, incubated for 5 min at 37°C, then 5 min on ice, then cells were passed through 70 μm filter for single cells suspension and downstream flow cytometric analysis.

#### Flow cytometry

Cells were preincubated with FcBlock to block non-specific Fc*g*R binding and Ghost Dye to detect dead cells. They were stained with antibodies targeting surface proteins for 30 min on ice. When needed, cells were then fixed and permeabilized for 30 min on ice and then stained with intracellular antibodies for 30 min on ice. All samples were acquired and analyzed with LSRII flow cytometer (Becton Dickson) and FlowJo software (TreeStar).

#### Cell sorting

After gating on morphology and singlets, CD45^+^Dead^-^ cells were gated as follows: Related to Figure 1: DC1 (Lin^-^F4/80^-^Ly6C^-^MHC-II^+^CD11c^+^Xcr1^+^CD11b^-^), DC2 (Lin^-^F4/80^-^Ly6C^-^MHC-II^+^CD11c^+^Xcr1^-^CD11b^+^), monocytes (Lin^-^F4/80^+^Ly6C^+^Xcr1^-^CD11b^+^), TAMs (Lin^-^F4/80^+^ Ly6C^-^MHC-II^+^Xcr1^-^ excluding CD11b^high^), MTMs (Lin^-^F4/80^+^Ly6C^-^MHC-II^+^Xcr1^-^CD11b^high^) (Lin = SiglecF, B220, Ly6G, TCR*b*, NK1.1). Each biological replicate was pooled tumors from PyMT mice, either two mice for DCs, or one mouse for monocytes, TAMs and MTMs, n = 2 for DC1, DC2, n = 3 for monocytes, MTMs and TAMs. Related to Figure 3: TAMs (F4/80^+^SiglecF^-^B220^-^ Ly6G^-^Ly6C^-^TCR*b*^-^NK1.1^-^Xcr1^-^ excluding CD11b^high^ [MTMs]) were sorted from tumors of *Irf8*^fl/fl^PyMT and CD11c^Cre^*Irf8*^fl/fl^PyMT mice. Each biological replicate was pooled tumors of one mouse, n = 2 for each group. Related to Figure 5: TCRβ^+^CD8α^+^ cells were sorted from tumors of *Irf8*^fl/fl^PyMT and MafB^iCre^*Irf8*^fl/fl^PyMT mice (n=1 per genotype). Related to Figure 6: Monocytes (Lin^-^ CD14^+^HLA-DR^low to neg^) and TAMs (Lin^-^CD14^+^HLA-DR^high^) were sorted from human RCC tumors (Lin = CD3, CD19, C56, CD15). Cell sorting was conducted on Aria II (Becton Dickson).

#### Bone marrow chimeras

Recipient mice (CD45.1.2^+^ S100a8^Cre^*Rosa26*^LSL-USA^PyMT) were irradiated with 900 Gyz at 8-12 weeks of age. Eighteen hours after irradiation, bone marrow was harvested from CD45.1^+^ C57Bl/6 and CD45.2^+^ CD11c^Cre^*Irf8*^fl/fl^ mice, mixed 1:1 and injected intravenously into irradiated recipient mice. Mice were maintained on sulfatrim antibiotic diet and aged to allow bone marrow to graft and PyMT tumors to grow. Upon large tumor burden (2000-3000 mm^3^), mice were euthanized and immune cells from tumor and draining lymph node were harvested for downstream analyses.

#### *In vitro* co-culture

Lymph nodes were harvested from OT-1 TCR-transgenic mice. Single-cell suspension of lymph nodes was obtained by homogenization between frosted ends of two histological slides and passed through 70 μm filters. Naïve CD8^+^ T cells were isolated using negative selection with magnetic beads, as per kit’s instructions. T cells were stained with CellTrace Violet and plated in 96 U-bottom wells (15,000 T cells/well), cocultured with various ratios of WT or KO TAMs sorted from tumors or migratory DC1 sorted from tumor-draining lymph nodes of chimeric mice in T cell media (RPMI supplemented with 10% FCS, 1 mM sodium pyruvate, non-essential amino acids [Gibco], 10 mM Hepes, 55 μM 2-Mercaptoethanol, 100 U/mL Penicillin G and 0.1 mg/ml Streptomycin) for 72 h at 37°C. Cells were then analyzed by flow cytometry.

#### RNA sequencing

For bulk RNA sequencing, immune populations were sorted into Trizol LS and flash frozen. RNA was extracted with chloroform. Isopropanol and linear acrylamide were added, and the RNA was precipitated with 75% ethanol. 0.649-1 ng total RNA with RNA integrity numbers 6.8-10 underwent amplification using SMART-Seq v4 Ultra Low Input RNA Kit (Clontech #63488). 15 ng amplified cDNA was used to prepare libraries with KAPA Hyper Prep Kit (Kapa Biosystems KK8504) using 8 cycles of PCR. Samples were barcoded and run on a HiSeq 2500 in 50bp/50bp paired end run using HiSeq3000/4000 SBS Kit (Illumina) for 30-40 million paired reads. The raw sequencing FASTQ files were aligned against the hg19 and mm10 assembly by STAR for human and mouse samples respectively. FASTQ files of TCGA ccRCC cohort (KIRC) were downloaded from GDC and processed by the same procedures. Gene level count values were then computed by the summarizeOverlaps function from the R package “GenomicAlignments” with UCSC hg19 or mm10 KnownGene as the base gene model for human and mouse samples respectively. The Union counting mode was used and only mapped paired reads were considered. FPKM (Fragments Per Kilobase Million) values were then computed from gene level counts by using fpkm function from the R package “DESeq2”.

For single cell RNA sequencing, around 10,000 FACS sorted live cells (1,000 cells/ul suspension) were suspended in 1X PBS (calcium and magnesium free) containing 0.04% BSA (A2153, SIGMA). Cell suspension was used as input in 10x chromium controller system (10x Genomics Inc., product code 120223) and cell barcoding was performed by gel beads in emulsion (GEMs) in assembly chip. GEM RT reaction was performed in thermocycler (53°C for 45 min, 85°C for 5 min, 4°C hold overnight). SPRIselect dynabeads (B23318, Beckman Coulter) were used for GEM recovery post RT. 2-50ng of DNA was used for target enrichment. cDNA amplification, fragmentation, end repair and A tailing preparation was performed as per manual instructions. High sensitivity DNA chips and Agilent 2100 bioanalyzer (Agilent Technologies) was used for gene expression library profile quality control and quantification at several steps. Quality control was performed twice before sequencing. Two barcoded scRNA samples were pooled together before sequencing. scRNA libraries were sequenced on NovaSeq 6000 S1 with sequencing depth of approximately 300 million reads.

### Quantification and Statistical Analysis

Figure 1B and Figure S3D: the regularized log transformation values generated by “DESeq2” of the top 500 varied genes were used for Principal Component Analysis and the first 2 principal components were shown in the plot. Analysis was performed on 17 samples: WT TAMs (n=3), MTMs (n=3), monocytes (n=3), DC1 (n=2), DC2 (n=2), and two pairs of WT and KO TAMs (n=2 each). All groups excluding WT and KO TAM pairs are shown in Figure 1B, all 17 samples are shown in Figure S3D.

Related to Figure 1C and Table S1, pairwise differentially expressed gene (DEG) analyses were done among 3 mouse sample groups, including monocytes, MTMs, and TAMs, by the R package DESeq2. Transcription factor genes (as annotated) were included if at least one pairwise comparison passed the following filters: FDR < 0.05, mean expression > 100, log_2_ fold change > 0.5, < −0.5. Related to Figure 3 and Figure S3, genes were significantly differentially expressed if passing the following filters: FDR < 0.05, mean expression > 50, log_2_ fold change > 2 or < −2. Mouse IRF8 WT-UP or IRF8 KO-UP signatures were built from DEG genes that passed filters listed above as well as mean FPKM > 1 and variance > 3. The mean FPKM value of selected gene was then calculated for each sample group. The log2 transformed mean FPKM values of signature genes were used for the empirical cumulative distribution plot by the R package “ggplot2”. An offset value of 0.001 was added to mean FPKM values prior to log2 transformation to avoid log2(0). DEG results for WT versus IRF8-KO TAMs (Figure S3C) were used in pathway analysis using IPA (QIAGEN Inc.) with the Ingenuity Knowledge Base as the reference set(Kramer et al., 2014). IPA evaluated the expression pattern of genes in canonical pathways and predicted the canonical pathways to be active or inactive.

The mouse DEG genes resulting from comparing WT to KO sample group with mean expression > 50, log2 fold change > 2 or < −2, FDR value < 0.05 underwent human homologous gene search. The corresponding human homologous genes were identified by referencing the Mouse Genome Informatics (MGI, http://www.informatics.jax.org/) database and NCBI Orthologs database (ftp://ftp.ncbi.nlm.nih.gov/gene/DATA/gene_orthologs.gz), and any gene without a human homolog was excluded from the human IRF8 signature. Gene Set Enrichment Analysis (GSEA) was done by applying the IRF8-activated human homologous genes against the DEG genes from human TAMs to tumor infiltrated human monocytes comparison through the R package “clusterProfiler” (Figure 6D). To ensure decent expression level in human cells, mouse IRF8-Activated genes were further filtered by having mean FPKM > 1 and variance > 3 among sorted human tumor monocytes and TAMs. These were used to build the human IRF8 signature (Figure 7A). To assess the IRF8 signature enrichment in human immune cell populations, Single-Sample Gene Set Enrichment Analysis (ssGSEA) was done by applying the human IRF8 signatures against the FPKM expression values of sorted human immune cell populations through the R package “GSVA” which estimates the IRF8 signature enrichment score for each of RNASeq samples (Figure 7B).

Along with IRF8 signature, the ssGSEA enrichment score of a CD8 T cells signature(Bindea et al., 2013) was also calculated for samples in TCGA KIRC RNASeq cohort. The KIRC cohort sample was classified as CD8^high^ when the CD8 T cells ssGSEA enrichment score is greater or equal to the median score and as CD8^low^ when it is smaller than the median score. Survival analyses and curves were performed and generated according to the Kaplan– Meier method using SAS v.9.4 (SAS Institute Inc., Cary, North Carolina). The enrichment scores for both the CD8 T cell signature and IRF8 signature were displayed along with median values, and T cell exhaustion (signature listed below) high or low based on median is color coded (Figure 7C). The log-rank test was used to determine statistical significance of the overall survival (OS) distributions between CD8^high^ and CD8^low^ groups in TCGA KIRC cohort (Figure 7D).

The KIRC cohort sample was classified as IRF8^high^ when the IRF8 signature ssGSEA enrichment score is greater or equal to the median score and as IRF8^low^ when it is smaller than the median score. DEG analysis between IRF8^high^ and ^low^ groups was performed and the results were subjected to GSEA analysis against 2 gene signatures, the Mac_CSF1 signature(Beck et al., 2009) and the T cell exhaustion/Checkpoint signature including several well-known T cell exhaustion markers and checkpoint markers, such as CTLA4, LAG3, PDCD1, CD274, PDCD1LG2, HAVCR2, BTLA and TIGIT (Figure 7E and 7F). Among the KIRC samples which have higher CD8 T cells score (CD8^high^ group), the survival analysis was done between IRF8^high^ and IRF8^low^ sample groups. The Kaplan–Meier curve and the log-rank test were generated using SAS v.9.4 (Figure 7G).

For scRNAseq analysis, the raw reads of single cell RNA sequencing from FASTQ files were aligned against the mm10 mouse genome and the UMI count of individual genes were captured by Cell Ranger v3.0.2 (10X Genomics) with “—expected-cells=1000” parameter setting and default value for th rest of required parameters. The R package Seurat 3.1.1(Butler et al., 2018; Stuart et al., 2019) was used for further QC, clustering, plotting and differentially expressed genes analyses. Based on the observation of low mitochondria read percentage distribution and PCA analysis results, cells filtering and batch correction procedure was not performed after merging cells from KO and wild-type mice. The SCTransform() function which implemented the sctransform package (https://github.com/ChristophH/sctransform) was used for UMI count normalization, scaling, and variant gene searching for further dimensionality reduction via calling RunPCA() function. The top 16 principal components (PCs) were chosen and used for Louvain clustering analysis using the FindClusters() function at resolution 0.6 after constructing the Shared Nearest Neighbor (SNN) Graph through calling FindNeighbors() on the selected PCs. A two-dimensional embedding of the data was generated using UMAP with the top 16 PCs as input by the RunUMAP() function. This first run of clustering revealed 11 cell clusters including T cell, innate-like T cell (ILTC), CD68+, possibly degraded T cell, and non-immune cell clusters. After excluding cells from CD68+, possibly degraded T cell, and non-immune cell clusters, a second run of clustering was performed on a total of 375 cells from WT tumors and 1525 cells from KO tumors, with top 11 PCs being selected after dimensionality reduction and 9 clusters found, including naïve CD8 (C1), effector/memory CD8 (C2a, C2b), exhaustion CD8 (C3a, C3b), transitioning CD8 (C4), ILTC (C5a, C5b), and proliferating CD8 (C6) clusters. Violin plots of selected genes were built by the VlnPlot() function. The Fisher’s Exact test was employed to test for the statistical significance of observed cell abundance difference of each cluster between KO and wild type groups. Differential gene expression analysis was conducted through the FindMarkers() function by using the Wilcoxon rank-sum test as the statistical test and a minimum of 10% cell expressing the gene and with FDR P < 0.05. Significantly differentially expressed genes (DEGs) between each set of paired clusters (a & b) were identified and enriched by immediate early genes (IEGs). The top most significant 40 genes between paired clusters were included for building the expression heatmaps by the DoHeatmap() function. Differential gene expression analysis was also conducted for each cluster by comparing the cluster cells to the cells from the other clusters as the reference cells. For those paired clusters, cells from their paired mate cluster were excluded from the reference cell group and the top 20 common DEGs among paired clusters were selected in building the cluster gene expression heatmap along with the top 20 DEGs from singleton clusters.

All data are displayed as mean +/− SEM. For pair-wise comparisons, unpaired or paired student t test, two-tailed was conducted using GraphPad Prism software. For tumor growth, 2-way ANOVA was performed using GraphPad Prism software. The analyzed data is provided in Tables S1-4.

### Data and Code Availability

The datasets supporting the current study are available from the corresponding author on request.

### Key Resources Table

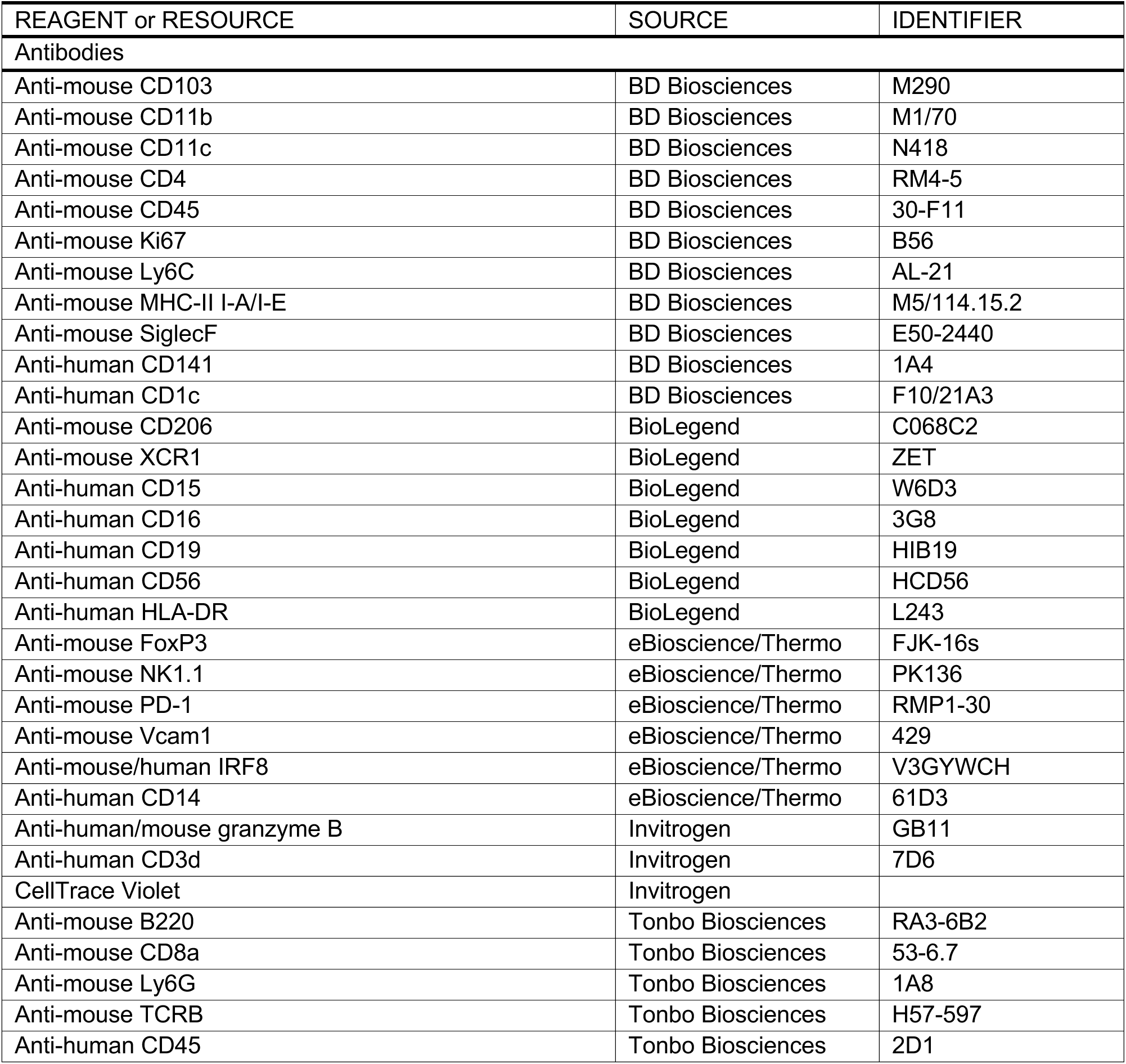
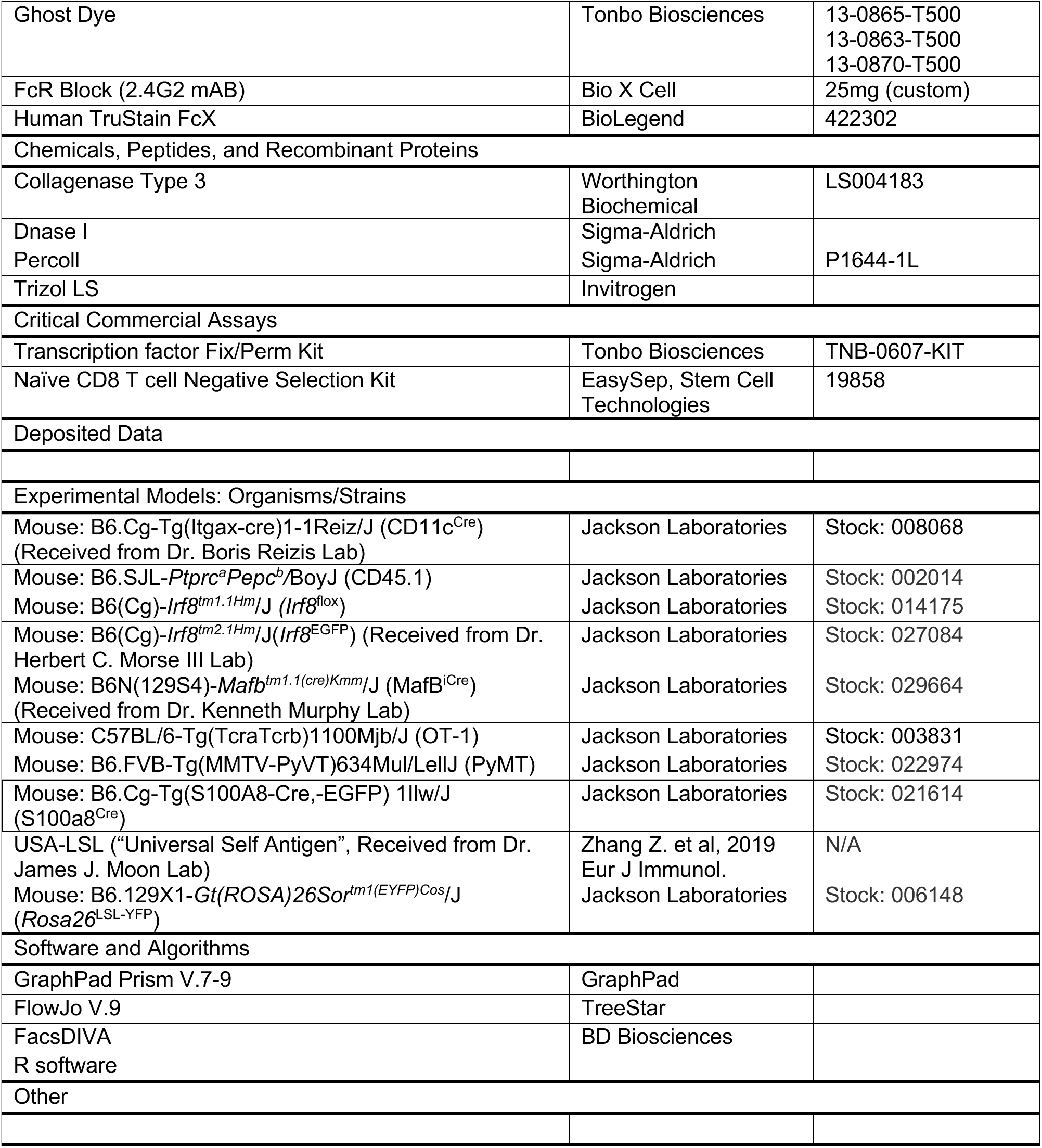

## Supplemental Information Titles and Legends

**Figure S1.**
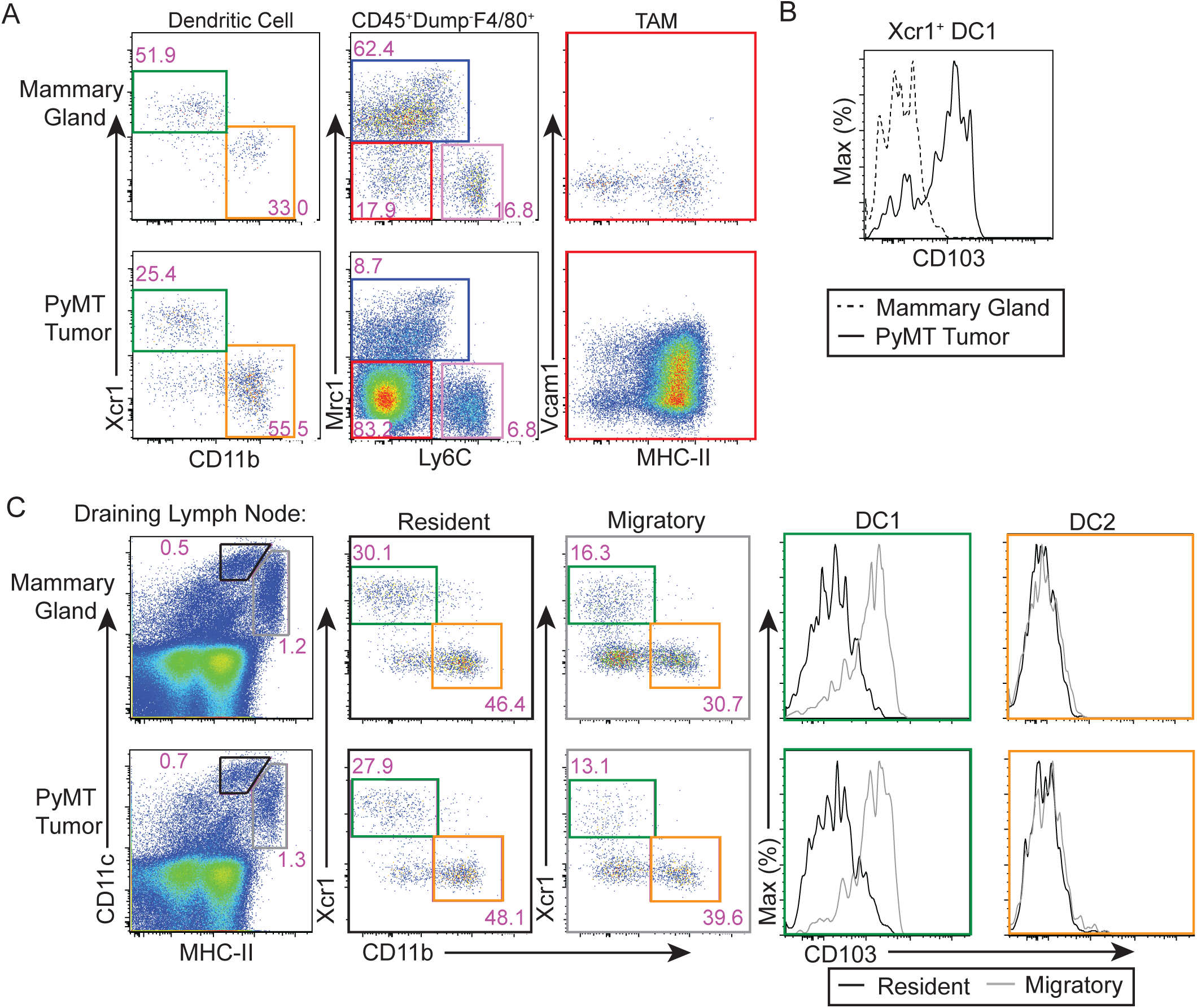
Mononuclear phagocytic cells of the mammary gland, tumor and their draining lymph nodes. (A) Flow cytometric gating strategy for mononuclear phagocytes in mammary gland and PyMT tumor: DC1s (green, CD45^+^Lin^-^F4/80^-^Ly6C^-^CD11c^+^MHC-II^+^Xcr1^+^), DC2s (orange, CD45^+^Lin^-^ F4/80^-^Ly6C^-^CD11c^+^MHC-II^+^CD11b^+^), monocytes (pink, CD45^+^Lin^-^F4/80^+^Ly6C^+^CD11b^+^), MTMs (blue, CD45^+^Lin^-^F4/80^+^Ly6C^-^Mrc1^+^) and TAMs (red, CD45^+^Lin^-^F4/80^+^Ly6C^-^Mrc1^-^) (Lin = Ly6G, B220, SiglecF, dead cells). Expression of Vcam1 and MHC-II on TAMs is also displayed. (B) CD103 expression on Xcr1^+^ DC1s from mammary gland (dashed line) and tumor (solid line). (C) Representative flow cytometric plots for resident (black, CD11c^high^MHCII^int^) and migratory (gray, CD11c^+^MHC-II^high^) DC1s (XCR1^+^) and DC2s (CD11b^+^) in mammary gland- or tumor-draining lymph node. Expression of CD103 in lymph node DC populations is shown for non-tumor bearing and tumor bearing mice. Data is representative of at least 3 independent experiments.

**Figure S2.**
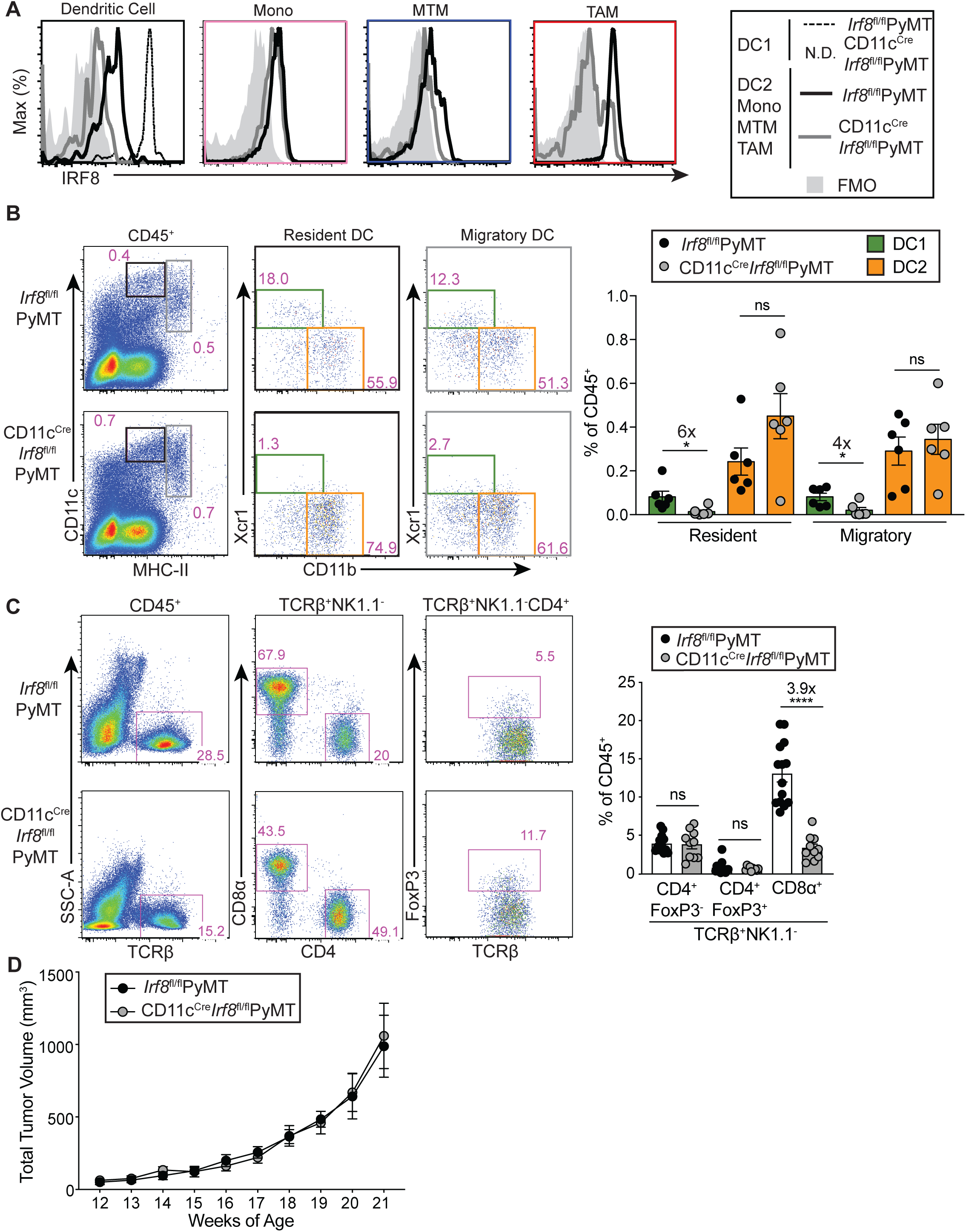
Leukocyte populations in tumor and tumor-draining lymph nodes of CD11c^Cre^*Irf8*^fl/fl^PyMT mice. (A) IRF8 protein expression as determined by flow cytometric analysis in DC1s, DC2s, monocytes, MTMs and TAMs from tumors in *Irf8*^fl/fl^PyMT (black line) and CD11c^Cre^*Irf8*^fl/fl^PyMT (gray line) mice. N.D.=no cells detected, WT DC1 (dashed line), and fluorescence minus one (FMO) sample unstained for IRF8. (B) Representative flow cytometric gating and quantification of tumor-draining lymph nodes in *Irf8*^fl/fl^PyMT (black circles) and CD11c^Cre^*Irf8*^fl/fl^PyMT (gray circles) mice (n=6 per group), resident DC (black box), migratory DC (gray box), DC1 (green bar) and DC2 (orange bar). (C) Representative flow plots and quantification of tumor-infiltrating T lymphocyte populations in *Irf8*^fl/fl^PyMT and CD11c^Cre^*Irf8*^fl/fl^PyMT mice. Each circle represents one mouse, data displayed as mean +/− SEM, fold change value of DCs from *Irf8*^fl/fl^PyMT versus CD11c^Cre^*Irf8*^fl/fl^PyMT mice are displayed above significant comparisons (unpaired student’s t test, two-tailed, * = p<0.05, **** = p<0.0001). (D) Tumor growth curves from CD11c^Cre^*Irf8*^fl/fl^PyMT (n=11) and littermate and cagemate *Irf8*^fl/fl^PyMT controls (n=8) (2-way ANOVA, not significant).

**Figure S3.**
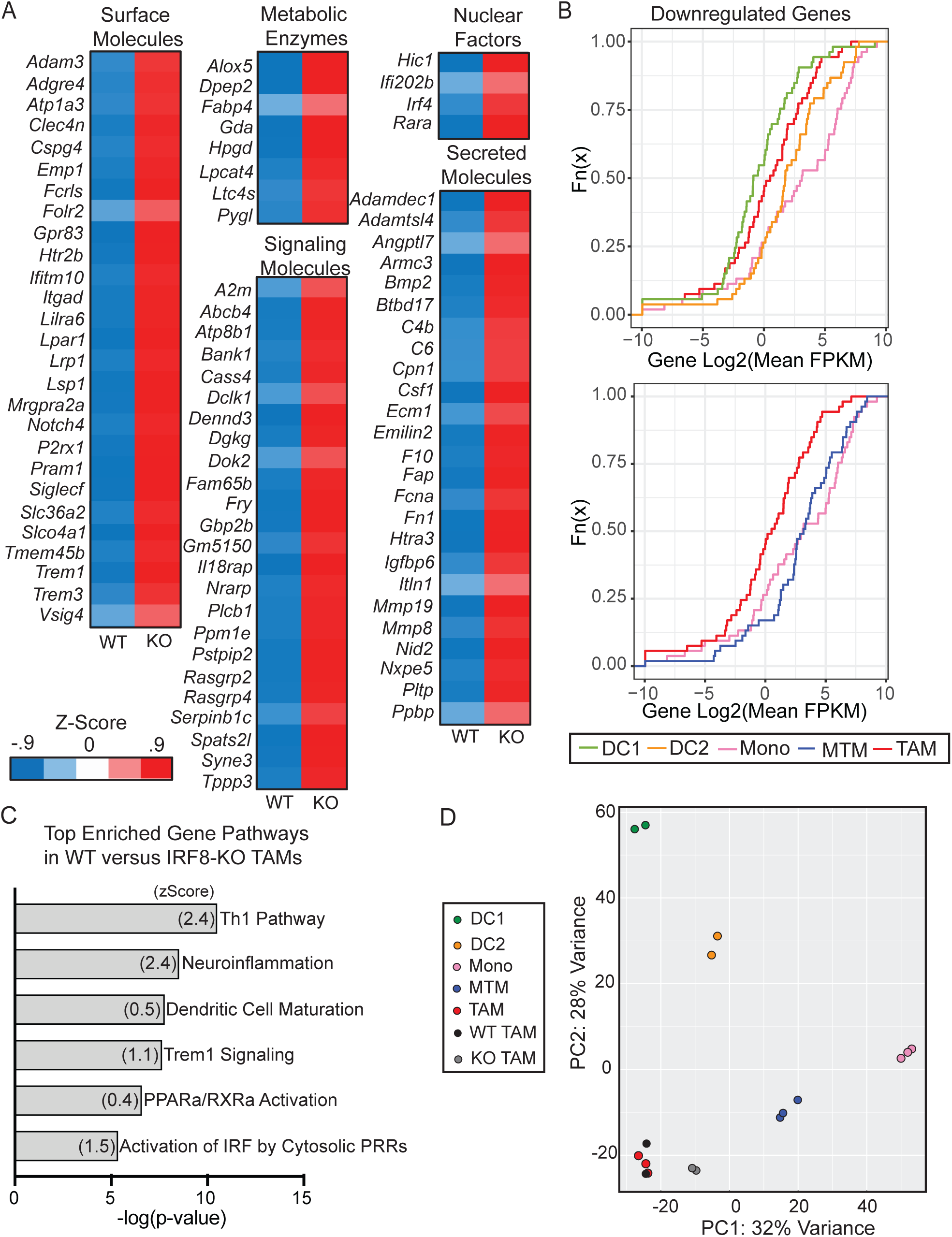
IRF8-dependent gene expression in TAMs. (A) Average z-score of genes significantly upregulated in TAMs sorted from tumors of *Irf8*^fl/fl^PyMT (wild-type, WT) and CD11c^Cre^*Irf8*^fl/fl^PyMT (knock-out, KO) mice. RNA was extracted and sequenced. Genes were grouped based on cell localization and function, excluding genes with unknown functions, pseudogenes and noncoding RNAs (*Art2a-ps, Nhsl2, Tmem181c-ps,* and *4933400F21Rik*) (base mean > 50, FDR < 0.05, log_2_ fold change > 2). (B) CDF plot displaying enrichment of IRF8-repressed gene signatures in PyMT tumor DC1 (green), DC2 (orange), monocyte (pink), MTM (blue), and TAM (red) RNAseq datasets from Figure 1. (C) Ingenuity Pathway Analysis among differentially expressed genes between WT and IRF8-KO TAMs. Most significantly enriched pathways in WT TAMs are shown. (D) Principal component analysis performed in Figure 1B, now displaying two WT and KO TAM pairs along with 5 groups of mononuclear phagocytes from PyMT tumors.

**Figure S4.**
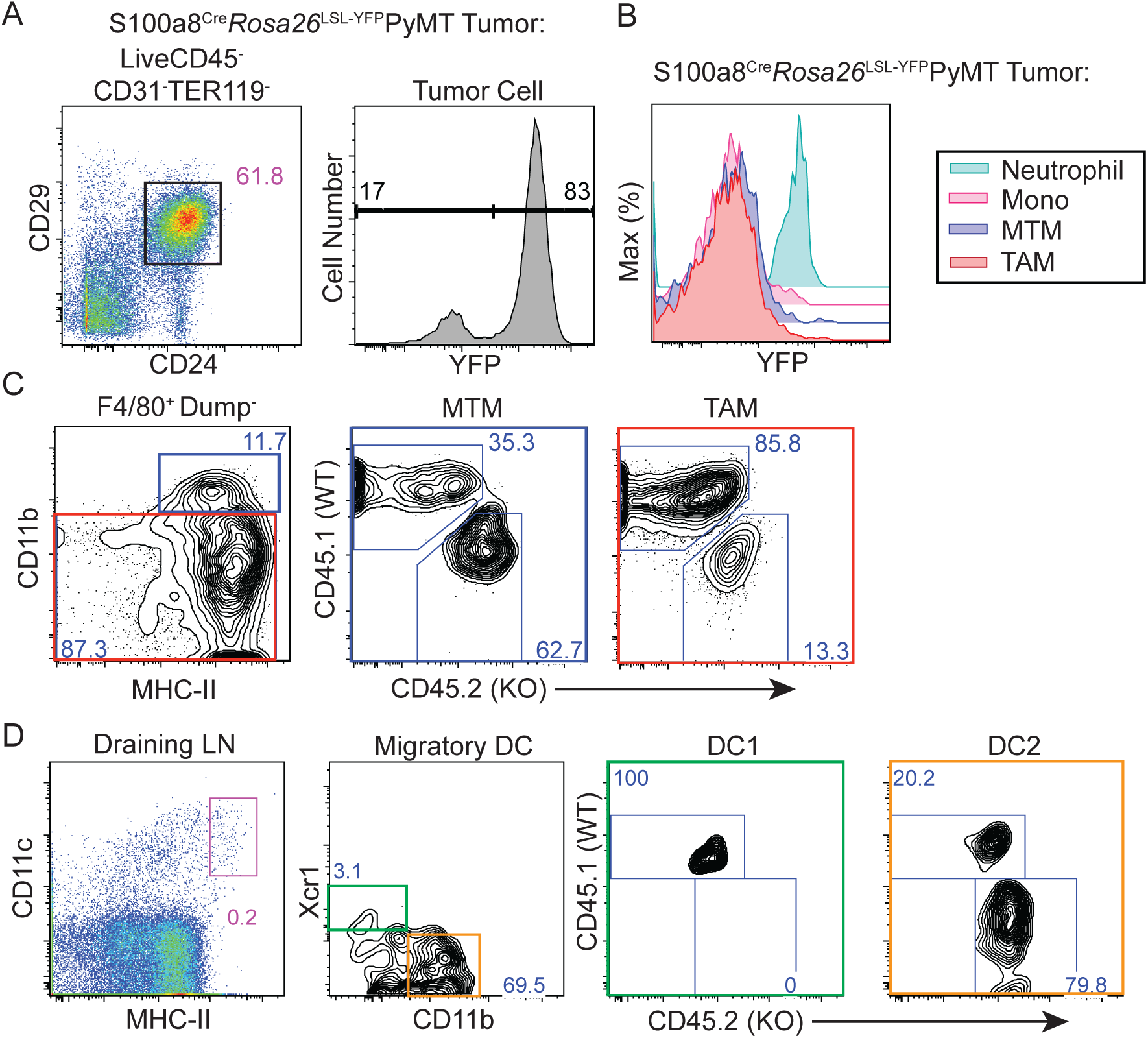
Characterization of S100a8^Cre^ in PyMT mice and chimeric mouse gating strategy. (A) Reporter for Cre activity (YFP expression) in PyMT tumor cells (CD45^-^CD31^-^TER119^-^ CD24^+^CD29^+^) of S100a8^Cre^*Rosa26*^LSL-YFP^PyMT mice. (B) Reporter for Cre activity (YFP expression) in tumor-infiltrating immune cells, including TAMs (red), MTMs (dark blue), monocytes (pink), and neutrophils (light blue, CD45^+^Ly6G^+^Ly6C^+^) of S100a8^Cre^*Rosa26*^LSL-YFP^PyMT mice. (C) Representative flow cytometric gating of tumor immune infiltrate in chimeric mice, related to Figure 3. Representative of nine independent experiments. (D) Representative flow cytometric gating of tumor-draining lymph node migratory DC1s from chimeric mice, related to Figure 4.

**Figure S5.**
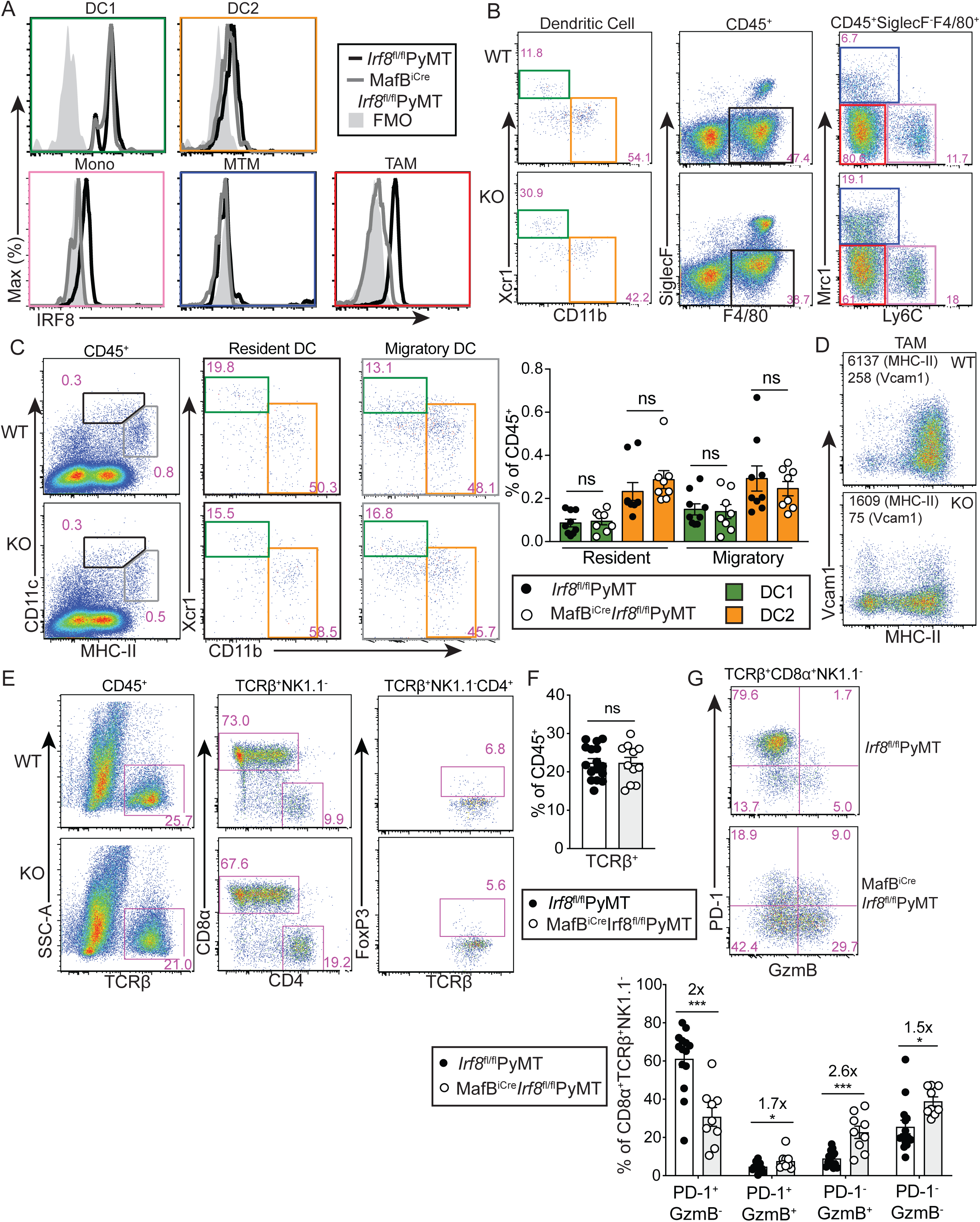
MafB^iCre^*Irf8*^fl/fl^PyMT mice display TAM defects while maintaining normal DC and T cell abundances. (A) IRF8 protein expression as determined by flow cytometric analysis in DC1s, DC2s, monocytes, MTMs and TAMs from tumors in *Irf8*^fl/fl^PyMT (black) and MafB^iCre^*Irf8*^fl/fl^PyMT (gray) mice, and a fluorescence minus one (FMO) sample unstained for IRF8. (B) Representative flow cytometric gating of DC1s, DC2s, F4/80^+^SiglecF^-^ cells, monocytes, MTMs and TAMs, corresponding to quantification displayed in Figure 5A. (C) Representative flow cytometric gating and quantification of resident (black) and migratory (gray) DC1s and DC2s in tumor-draining lymph nodes of *Irf8*^fl/fl^PyMT (wild-type, WT, black circles, n=9) and MafB^iCre^*Irf8*^fl/fl^PyMT (knock-out, KO, white circles, n=8) mice. (D) Representative flow cytometric plot and quantification of fold change in geometric mean fluorescence intensity (MFI) of MHC-II and Vcam1 relative to WT TAMs, MFI indicated on representative plot. (E) Representative flow quantification of tumor-infiltrating lymphocytes in WT and KO mice, corresponding to quantification displayed in Figure 5C. (F) Quantification of total TCRβ^+^ cells in tumors of *Irf8*^fl/fl^PyMT (black circle, n=16) and MafB^iCre^*Irf8*^fl/fl^PyMT (white circle, n=11) mice. (G) Representative flow cytometric analysis (top) and quantification (bottom) of PD-1 and granzyme B (GzmB) expression among TCRβ^+^CD8α^+^NK1.1^-^ cells of WT (n=14) and KO (n=9) mouse tumors. Data is displayed as mean +/− SEM, fold change value of WT versus KO means are displayed above significant comparisons (D, F, G: unpaired student’s t test, two-tailed, * p=<0.05, *** = p<0.001).

**Figure S6.**
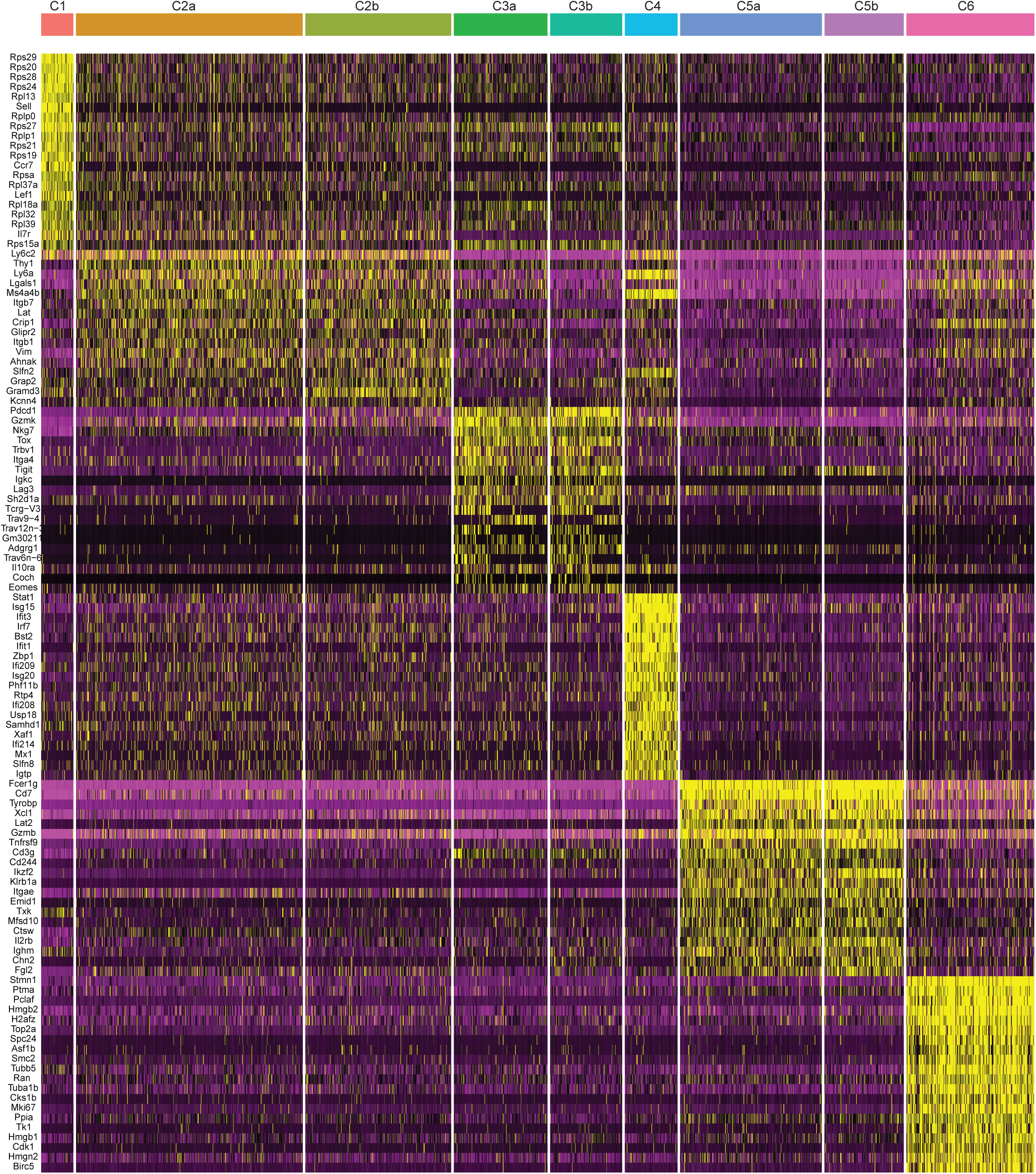
Differentially expressed transcripts among scRNAseq clusters. Expression of genes in all cells in all 9 clusters. Top twenty differentially expressed genes (DEGs) among each cluster relative to all the others are shown. For clusters with “a” and “b”, DEGs were determined by shared DEGs among “a” cluster versus all other clusters minus “b”, and “b” cluster versus all other clusters minus “a”.

**Figure S7.**
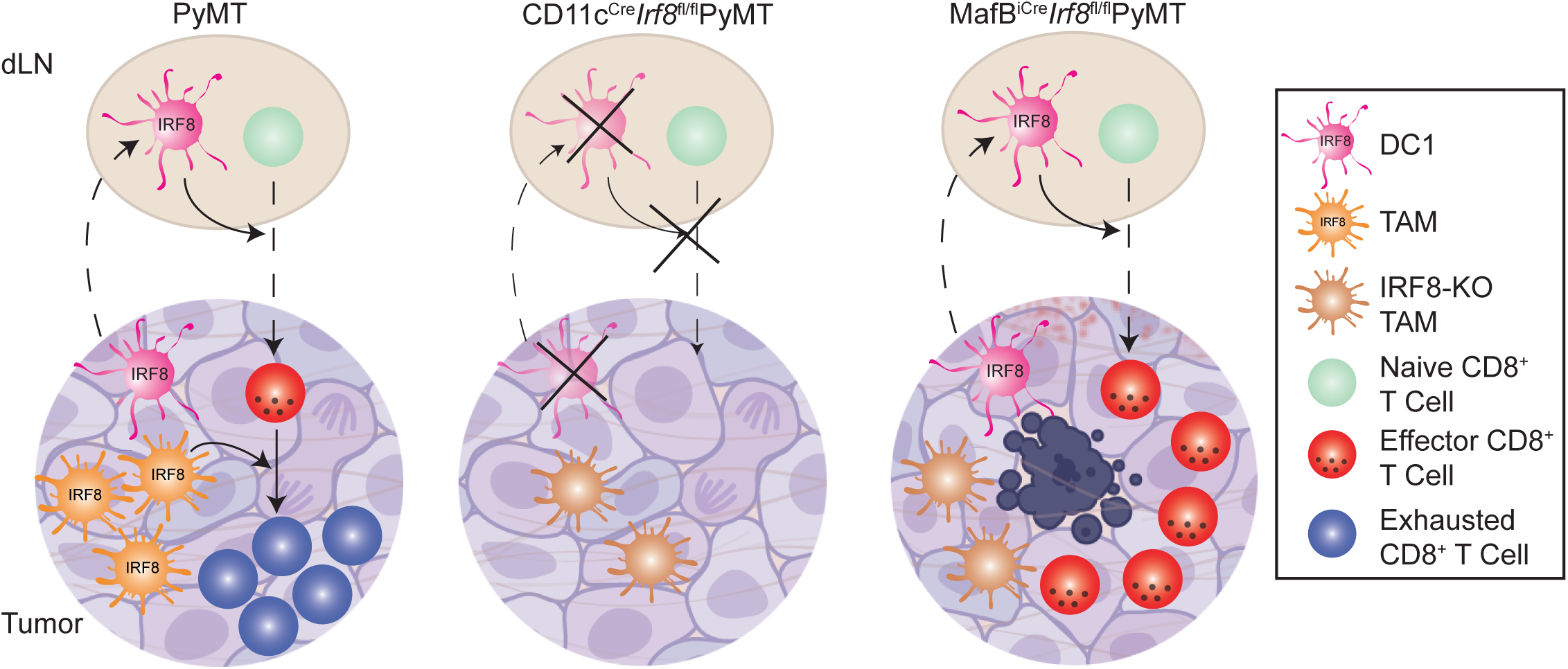
IRF8 governs TAM control of T cell exhaustion. Model summarizing findings, showing DC1, TAM and CD8^+^ T cell fates in tumor-draining lymph nodes (dLN) and tumor of PyMT, CD11c^Cre^*Irf8*^fl/fl^PyMT, and MafB^iCre^*Irf8*^fl/fl^PyMT mice.

## Supplemental Tables

**Table S1.**
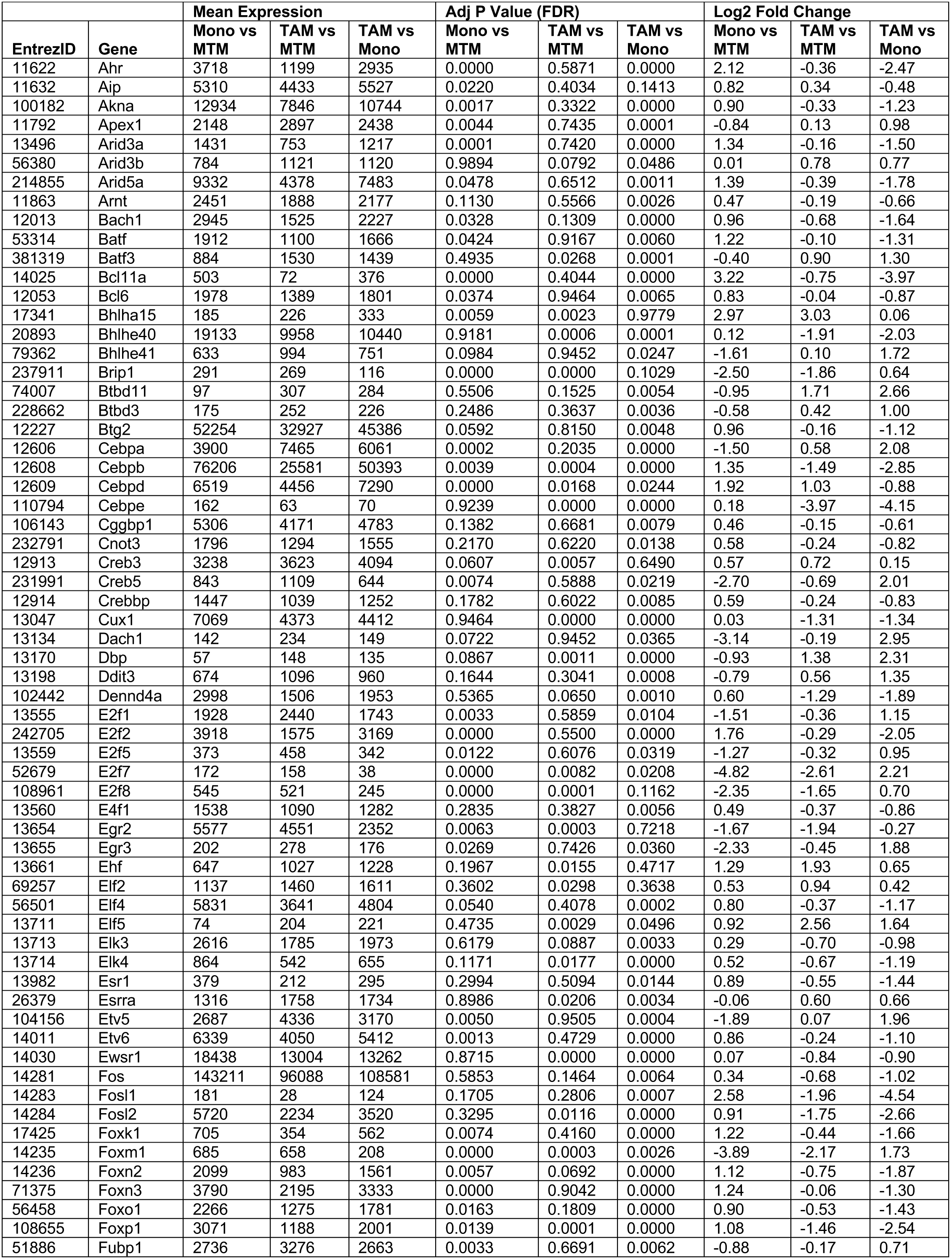
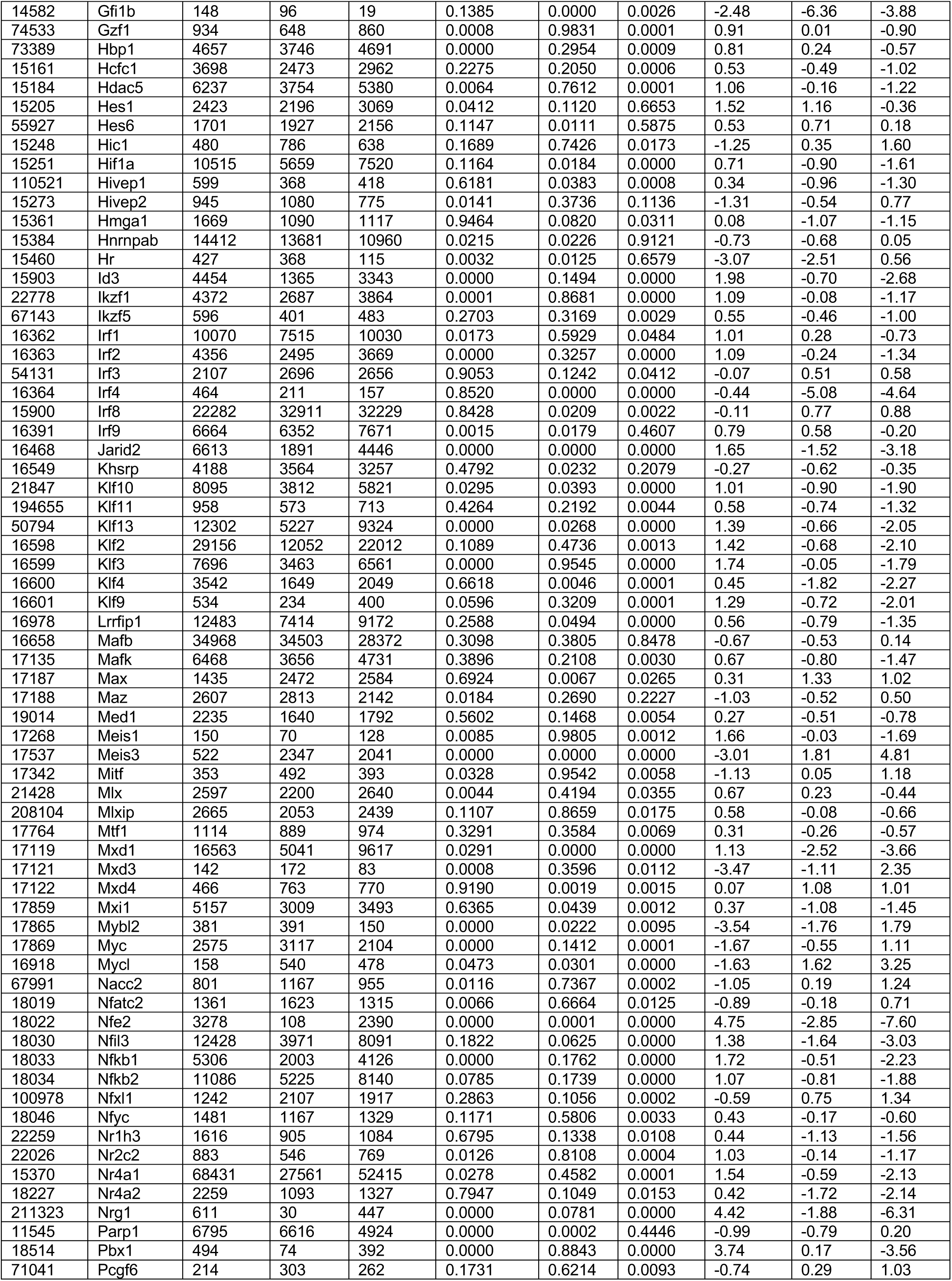
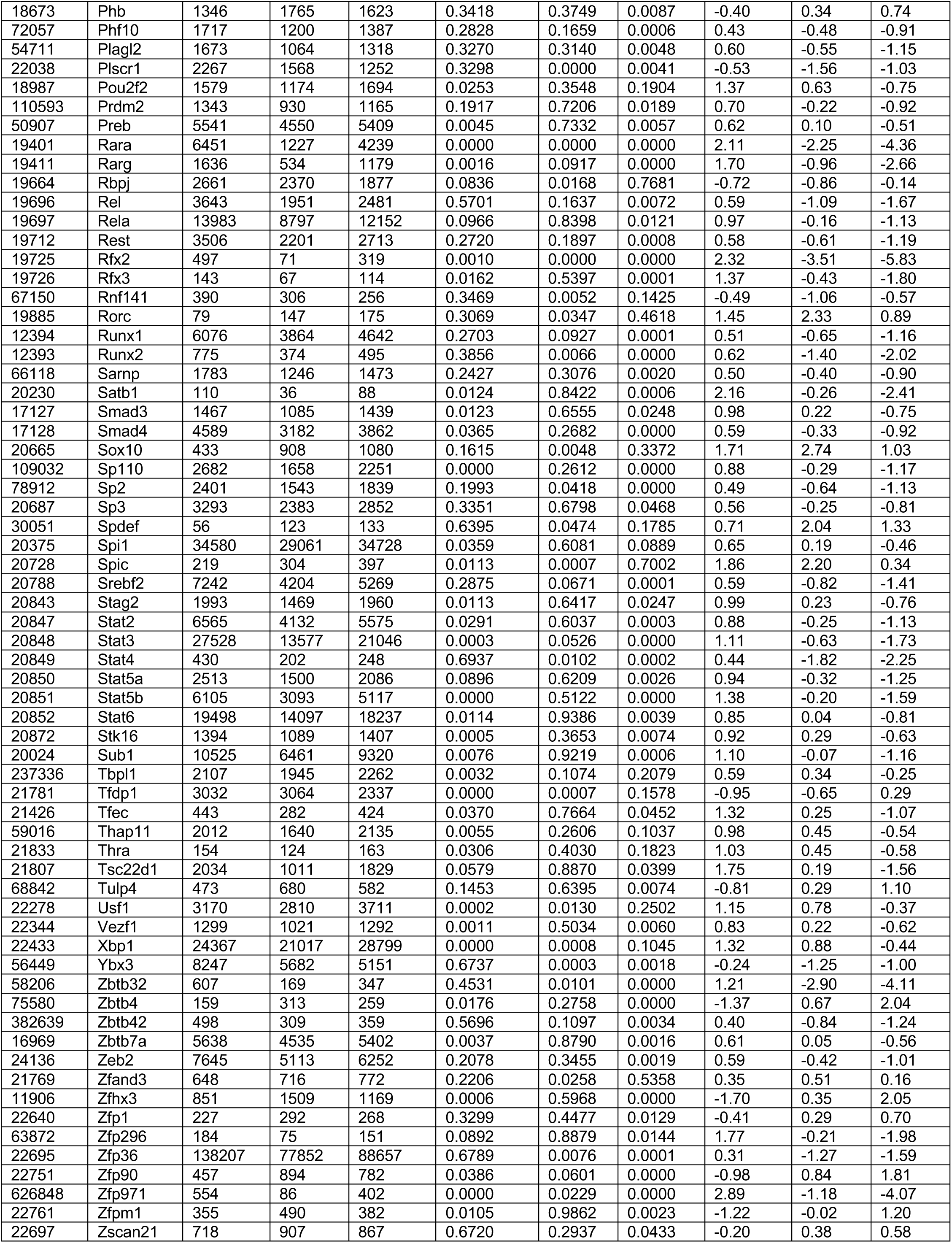
Differentially expressed transcription factor genes among monocytes (Mono), MTMs and TAMs.

|  |  | Mean Expression |  |  | Adj P Value (FDR) |  |  | Log2 Fold Change |  |  |
| --- | --- | --- | --- | --- | --- | --- | --- | --- | --- | --- |
| EntrezID | Gene | Mono vs MTM | TAM vs MTM | TAM vs Mono | Mono vs MTM | TAM vs MTM | TAM vs Mono | Mono vs MTM | TAM vs MTM | TAM vs Mono |
| 11622 | Ahr | 3718 | 1199 | 2935 | 0.0000 | 0.5871 | 0.0000 | 2.12 | -0.36 | -2.47 |
| 11632 | Aip | 5310 | 4433 | 5527 | 0.0220 | 0.4034 | 0.1413 | 0.82 | 0.34 | -0.48 |
| 100182 | Akna | 12934 | 7846 | 10744 | 0.0017 | 0.3322 | 0.0000 | 0.90 | -0.33 | -1.23 |
| 11792 | Apex1 | 2148 | 2897 | 2438 | 0.0044 | 0.7435 | 0.0001 | -0.84 | 0.13 | 0.98 |
| 13496 | Arid3a | 1431 | 753 | 1217 | 0.0001 | 0.7420 | 0.0000 | 1.34 | -0.16 | -1.50 |
| 56380 | Arid3b | 784 | 1121 | 1120 | 0.9894 | 0.0792 | 0.0486 | 0.01 | 0.78 | 0.77 |
| 214855 | Arid5a | 9332 | 4378 | 7483 | 0.0478 | 0.6512 | 0.0011 | 1.39 | -0.39 | -1.78 |
| 11863 | Arnt | 2451 | 1888 | 2177 | 0.1130 | 0.5566 | 0.0026 | 0.47 | -0.19 | -0.66 |
| 12013 | Bach1 | 2945 | 1525 | 2227 | 0.0328 | 0.1309 | 0.0000 | 0.96 | -0.68 | -1.64 |
| 53314 | Batf | 1912 | 1100 | 1666 | 0.0424 | 0.9167 | 0.0060 | 1.22 | -0.10 | -1.31 |
| 381319 | Batf3 | 884 | 1530 | 1439 | 0.4935 | 0.0268 | 0.0001 | -0.40 | 0.90 | 1.30 |
| 14025 | Bcl11a | 503 | 72 | 376 | 0.0000 | 0.4044 | 0.0000 | 3.22 | -0.75 | -3.97 |
| 12053 | Bcl6 | 1978 | 1389 | 1801 | 0.0374 | 0.9464 | 0.0065 | 0.83 | -0.04 | -0.87 |
| 17341 | Bhlha15 | 185 | 226 | 333 | 0.0059 | 0.0023 | 0.9779 | 2.97 | 3.03 | 0.06 |
| 20893 | Bhlhe40 | 19133 | 9958 | 10440 | 0.9181 | 0.0006 | 0.0001 | 0.12 | -1.91 | -2.03 |
| 79362 | Bhlhe41 | 633 | 994 | 751 | 0.0984 | 0.9452 | 0.0247 | -1.61 | 0.10 | 1.72 |
| 237911 | Brip1 | 291 | 269 | 116 | 0.0000 | 0.0000 | 0.1029 | -2.50 | -1.86 | 0.64 |
| 74007 | Btbd11 | 97 | 307 | 284 | 0.5506 | 0.1525 | 0.0054 | -0.95 | 1.71 | 2.66 |
| 228662 | Btbd3 | 175 | 252 | 226 | 0.2486 | 0.3637 | 0.0036 | -0.58 | 0.42 | 1.00 |
| 12227 | Btg2 | 52254 | 32927 | 45386 | 0.0592 | 0.8150 | 0.0048 | 0.96 | -0.16 | -1.12 |
| 12606 | Cebpa | 3900 | 7465 | 6061 | 0.0002 | 0.2035 | 0.0000 | -1.50 | 0.58 | 2.08 |
| 12608 | Cebpb | 76206 | 25581 | 50393 | 0.0039 | 0.0004 | 0.0000 | 1.35 | -1.49 | -2.85 |
| 12609 | Cebpd | 6519 | 4456 | 7290 | 0.0000 | 0.0168 | 0.0244 | 1.92 | 1.03 | -0.88 |
| 110794 | Cebpe | 162 | 63 | 70 | 0.9239 | 0.0000 | 0.0000 | 0.18 | -3.97 | -4.15 |
| 106143 | Cggbp1 | 5306 | 4171 | 4783 | 0.1382 | 0.6681 | 0.0079 | 0.46 | -0.15 | -0.61 |
| 232791 | Cnot3 | 1796 | 1294 | 1555 | 0.2170 | 0.6220 | 0.0138 | 0.58 | -0.24 | -0.82 |
| 12913 | Creb3 | 3238 | 3623 | 4094 | 0.0607 | 0.0057 | 0.6490 | 0.57 | 0.72 | 0.15 |
| 231991 | Creb5 | 843 | 1109 | 644 | 0.0074 | 0.5888 | 0.0219 | -2.70 | -0.69 | 2.01 |
| 12914 | Crebbp | 1447 | 1039 | 1252 | 0.1782 | 0.6022 | 0.0085 | 0.59 | -0.24 | -0.83 |
| 13047 | Cux1 | 7069 | 4373 | 4412 | 0.9464 | 0.0000 | 0.0000 | 0.03 | -1.31 | -1.34 |
| 13134 | Dach1 | 142 | 234 | 149 | 0.0722 | 0.9452 | 0.0365 | -3.14 | -0.19 | 2.95 |
| 13170 | Dbp | 57 | 148 | 135 | 0.0867 | 0.0011 | 0.0000 | -0.93 | 1.38 | 2.31 |
| 13198 | Ddit3 | 674 | 1096 | 960 | 0.1644 | 0.3041 | 0.0008 | -0.79 | 0.56 | 1.35 |
| 102442 | Dennd4a | 2998 | 1506 | 1953 | 0.5365 | 0.0650 | 0.0010 | 0.60 | -1.29 | -1.89 |
| 13555 | E2f1 | 1928 | 2440 | 1743 | 0.0033 | 0.5859 | 0.0104 | -1.51 | -0.36 | 1.15 |
| 242705 | E2f2 | 3918 | 1575 | 3169 | 0.0000 | 0.5500 | 0.0000 | 1.76 | -0.29 | -2.05 |
| 13559 | E2f5 | 373 | 458 | 342 | 0.0122 | 0.6076 | 0.0319 | -1.27 | -0.32 | 0.95 |
| 52679 | E2f7 | 172 | 158 | 38 | 0.0000 | 0.0082 | 0.0208 | -4.82 | -2.61 | 2.21 |
| 108961 | E2f8 | 545 | 521 | 245 | 0.0000 | 0.0001 | 0.1162 | -2.35 | -1.65 | 0.70 |
| 13560 | E4f1 | 1538 | 1090 | 1282 | 0.2835 | 0.3827 | 0.0056 | 0.49 | -0.37 | -0.86 |
| 13654 | Egr2 | 5577 | 4551 | 2352 | 0.0063 | 0.0003 | 0.7218 | -1.67 | -1.94 | -0.27 |
| 13655 | Egr3 | 202 | 278 | 176 | 0.0269 | 0.7426 | 0.0360 | -2.33 | -0.45 | 1.88 |
| 13661 | Ehf | 647 | 1027 | 1228 | 0.1967 | 0.0155 | 0.4717 | 1.29 | 1.93 | 0.65 |
| 69257 | Elf2 | 1137 | 1460 | 1611 | 0.3602 | 0.0298 | 0.3638 | 0.53 | 0.94 | 0.42 |
| 56501 | Elf4 | 5831 | 3641 | 4804 | 0.0540 | 0.4078 | 0.0002 | 0.80 | -0.37 | -1.17 |
| 13711 | Elf5 | 74 | 204 | 221 | 0.4735 | 0.0029 | 0.0496 | 0.92 | 2.56 | 1.64 |
| 13713 | Elk3 | 2616 | 1785 | 1973 | 0.6179 | 0.0887 | 0.0033 | 0.29 | -0.70 | -0.98 |
| 13714 | Elk4 | 864 | 542 | 655 | 0.1171 | 0.0177 | 0.0000 | 0.52 | -0.67 | -1.19 |
| 13982 | Esr1 | 379 | 212 | 295 | 0.2994 | 0.5094 | 0.0144 | 0.89 | -0.55 | -1.44 |
| 26379 | Esrra | 1316 | 1758 | 1734 | 0.8986 | 0.0206 | 0.0034 | -0.06 | 0.60 | 0.66 |
| 104156 | Etv5 | 2687 | 4336 | 3170 | 0.0050 | 0.9505 | 0.0004 | -1.89 | 0.07 | 1.96 |
| 14011 | Etv6 | 6339 | 4050 | 5412 | 0.0013 | 0.4729 | 0.0000 | 0.86 | -0.24 | -1.10 |
| 14030 | Ewsr1 | 18438 | 13004 | 13262 | 0.8715 | 0.0000 | 0.0000 | 0.07 | -0.84 | -0.90 |
| 14281 | Fos | 143211 | 96088 | 108581 | 0.5853 | 0.1464 | 0.0064 | 0.34 | -0.68 | -1.02 |
| 14283 | Fosl1 | 181 | 28 | 124 | 0.1705 | 0.2806 | 0.0007 | 2.58 | -1.96 | -4.54 |
| 14284 | Fosl2 | 5720 | 2234 | 3520 | 0.3295 | 0.0116 | 0.0000 | 0.91 | -1.75 | -2.66 |
| 17425 | Foxk1 | 705 | 354 | 562 | 0.0074 | 0.4160 | 0.0000 | 1.22 | -0.44 | -1.66 |
| 14235 | Foxm1 | 685 | 658 | 208 | 0.0000 | 0.0003 | 0.0026 | -3.89 | -2.17 | 1.73 |
| 14236 | Foxn2 | 2099 | 983 | 1561 | 0.0057 | 0.0692 | 0.0000 | 1.12 | -0.75 | -1.87 |
| 71375 | Foxn3 | 3790 | 2195 | 3333 | 0.0000 | 0.9042 | 0.0000 | 1.24 | -0.06 | -1.30 |
| 56458 | Foxo1 | 2266 | 1275 | 1781 | 0.0163 | 0.1809 | 0.0000 | 0.90 | -0.53 | -1.43 |
| 108655 | Foxp1 | 3071 | 1188 | 2001 | 0.0139 | 0.0001 | 0.0000 | 1.08 | -1.46 | -2.54 |
| 51886 | Fubp1 | 2736 | 3276 | 2663 | 0.0033 | 0.6691 | 0.0062 | -0.88 | -0.17 | 0.71 |
| 14582 | Gfi1b | 148 | 96 | 19 | 0.1385 | 0.0000 | 0.0026 | -2.48 | -6.36 | -3.88 |
| 74533 | Gzf1 | 934 | 648 | 860 | 0.0008 | 0.9831 | 0.0001 | 0.91 | 0.01 | -0.90 |
| 73389 | Hbp1 | 4657 | 3746 | 4691 | 0.0000 | 0.2954 | 0.0009 | 0.81 | 0.24 | -0.57 |
| 15161 | Hcfc1 | 3698 | 2473 | 2962 | 0.2275 | 0.2050 | 0.0006 | 0.53 | -0.49 | -1.02 |
| 15184 | Hdac5 | 6237 | 3754 | 5380 | 0.0064 | 0.7612 | 0.0001 | 1.06 | -0.16 | -1.22 |
| 15205 | Hes1 | 2423 | 2196 | 3069 | 0.0412 | 0.1120 | 0.6653 | 1.52 | 1.16 | -0.36 |
| 55927 | Hes6 | 1701 | 1927 | 2156 | 0.1147 | 0.0111 | 0.5875 | 0.53 | 0.71 | 0.18 |
| 15248 | Hic1 | 480 | 786 | 638 | 0.1689 | 0.7426 | 0.0173 | -1.25 | 0.35 | 1.60 |
| 15251 | Hif1a | 10515 | 5659 | 7520 | 0.1164 | 0.0184 | 0.0000 | 0.71 | -0.90 | -1.61 |
| 110521 | Hivep1 | 599 | 368 | 418 | 0.6181 | 0.0383 | 0.0008 | 0.34 | -0.96 | -1.30 |
| 15273 | Hivep2 | 945 | 1080 | 775 | 0.0141 | 0.3736 | 0.1136 | -1.31 | -0.54 | 0.77 |
| 15361 | Hmga1 | 1669 | 1090 | 1117 | 0.9464 | 0.0820 | 0.0311 | 0.08 | -1.07 | -1.15 |
| 15384 | Hnrnpab | 14412 | 13681 | 10960 | 0.0215 | 0.0226 | 0.9121 | -0.73 | -0.68 | 0.05 |
| 15460 | Hr | 427 | 368 | 115 | 0.0032 | 0.0125 | 0.6579 | -3.07 | -2.51 | 0.56 |
| 15903 | Id3 | 4454 | 1365 | 3343 | 0.0000 | 0.1494 | 0.0000 | 1.98 | -0.70 | -2.68 |
| 22778 | Ikzf1 | 4372 | 2687 | 3864 | 0.0001 | 0.8681 | 0.0000 | 1.09 | -0.08 | -1.17 |
| 67143 | Ikzf5 | 596 | 401 | 483 | 0.2703 | 0.3169 | 0.0029 | 0.55 | -0.46 | -1.00 |
| 16362 | Irf1 | 10070 | 7515 | 10030 | 0.0173 | 0.5929 | 0.0484 | 1.01 | 0.28 | -0.73 |
| 16363 | Irf2 | 4356 | 2495 | 3669 | 0.0000 | 0.3257 | 0.0000 | 1.09 | -0.24 | -1.34 |
| 54131 | Irf3 | 2107 | 2696 | 2656 | 0.9053 | 0.1242 | 0.0412 | -0.07 | 0.51 | 0.58 |
| 16364 | Irf4 | 464 | 211 | 157 | 0.8520 | 0.0000 | 0.0000 | -0.44 | -5.08 | -4.64 |
| 15900 | Irf8 | 22282 | 32911 | 32229 | 0.8428 | 0.0209 | 0.0022 | -0.11 | 0.77 | 0.88 |
| 16391 | Irf9 | 6664 | 6352 | 7671 | 0.0015 | 0.0179 | 0.4607 | 0.79 | 0.58 | -0.20 |
| 16468 | Jarid2 | 6613 | 1891 | 4446 | 0.0000 | 0.0000 | 0.0000 | 1.65 | -1.52 | -3.18 |
| 16549 | Khsrp | 4188 | 3564 | 3257 | 0.4792 | 0.0232 | 0.2079 | -0.27 | -0.62 | -0.35 |
| 21847 | Klf10 | 8095 | 3812 | 5821 | 0.0295 | 0.0393 | 0.0000 | 1.01 | -0.90 | -1.90 |
| 194655 | Klf11 | 958 | 573 | 713 | 0.4264 | 0.2192 | 0.0044 | 0.58 | -0.74 | -1.32 |
| 50794 | Klf13 | 12302 | 5227 | 9324 | 0.0000 | 0.0268 | 0.0000 | 1.39 | -0.66 | -2.05 |
| 16598 | Klf2 | 29156 | 12052 | 22012 | 0.1089 | 0.4736 | 0.0013 | 1.42 | -0.68 | -2.10 |
| 16599 | Klf3 | 7696 | 3463 | 6561 | 0.0000 | 0.9545 | 0.0000 | 1.74 | -0.05 | -1.79 |
| 16600 | Klf4 | 3542 | 1649 | 2049 | 0.6618 | 0.0046 | 0.0001 | 0.45 | -1.82 | -2.27 |
| 16601 | Klf9 | 534 | 234 | 400 | 0.0596 | 0.3209 | 0.0001 | 1.29 | -0.72 | -2.01 |
| 16978 | Lrrfip1 | 12483 | 7414 | 9172 | 0.2588 | 0.0494 | 0.0000 | 0.56 | -0.79 | -1.35 |
| 16658 | Mafb | 34968 | 34503 | 28372 | 0.3098 | 0.3805 | 0.8478 | -0.67 | -0.53 | 0.14 |
| 17135 | Mafk | 6468 | 3656 | 4731 | 0.3896 | 0.2108 | 0.0030 | 0.67 | -0.80 | -1.47 |
| 17187 | Max | 1435 | 2472 | 2584 | 0.6924 | 0.0067 | 0.0265 | 0.31 | 1.33 | 1.02 |
| 17188 | Maz | 2607 | 2813 | 2142 | 0.0184 | 0.2690 | 0.2227 | -1.03 | -0.52 | 0.50 |
| 19014 | Med1 | 2235 | 1640 | 1792 | 0.5602 | 0.1468 | 0.0054 | 0.27 | -0.51 | -0.78 |
| 17268 | Meis1 | 150 | 70 | 128 | 0.0085 | 0.9805 | 0.0012 | 1.66 | -0.03 | -1.69 |
| 17537 | Meis3 | 522 | 2347 | 2041 | 0.0000 | 0.0000 | 0.0000 | -3.01 | 1.81 | 4.81 |
| 17342 | Mitf | 353 | 492 | 393 | 0.0328 | 0.9542 | 0.0058 | -1.13 | 0.05 | 1.18 |
| 21428 | Mlx | 2597 | 2200 | 2640 | 0.0044 | 0.4194 | 0.0355 | 0.67 | 0.23 | -0.44 |
| 208104 | Mlxip | 2665 | 2053 | 2439 | 0.1107 | 0.8659 | 0.0175 | 0.58 | -0.08 | -0.66 |
| 17764 | Mtf1 | 1114 | 889 | 974 | 0.3291 | 0.3584 | 0.0069 | 0.31 | -0.26 | -0.57 |
| 17119 | Mxd1 | 16563 | 5041 | 9617 | 0.0291 | 0.0000 | 0.0000 | 1.13 | -2.52 | -3.66 |
| 17121 | Mxd3 | 142 | 172 | 83 | 0.0008 | 0.3596 | 0.0112 | -3.47 | -1.11 | 2.35 |
| 17122 | Mxd4 | 466 | 763 | 770 | 0.9190 | 0.0019 | 0.0015 | 0.07 | 1.08 | 1.01 |
| 17859 | Mxi1 | 5157 | 3009 | 3493 | 0.6365 | 0.0439 | 0.0012 | 0.37 | -1.08 | -1.45 |
| 17865 | Mybl2 | 381 | 391 | 150 | 0.0000 | 0.0222 | 0.0095 | -3.54 | -1.76 | 1.79 |
| 17869 | Myc | 2575 | 3117 | 2104 | 0.0000 | 0.1412 | 0.0001 | -1.67 | -0.55 | 1.11 |
| 16918 | Mycl | 158 | 540 | 478 | 0.0473 | 0.0301 | 0.0000 | -1.63 | 1.62 | 3.25 |
| 67991 | Nacc2 | 801 | 1167 | 955 | 0.0116 | 0.7367 | 0.0002 | -1.05 | 0.19 | 1.24 |
| 18019 | Nfatc2 | 1361 | 1623 | 1315 | 0.0066 | 0.6664 | 0.0125 | -0.89 | -0.18 | 0.71 |
| 18022 | Nfe2 | 3278 | 108 | 2390 | 0.0000 | 0.0001 | 0.0000 | 4.75 | -2.85 | -7.60 |
| 18030 | Nfil3 | 12428 | 3971 | 8091 | 0.1822 | 0.0625 | 0.0000 | 1.38 | -1.64 | -3.03 |
| 18033 | Nfkb1 | 5306 | 2003 | 4126 | 0.0000 | 0.1762 | 0.0000 | 1.72 | -0.51 | -2.23 |
| 18034 | Nfkb2 | 11086 | 5225 | 8140 | 0.0785 | 0.1739 | 0.0000 | 1.07 | -0.81 | -1.88 |
| 100978 | Nfxl1 | 1242 | 2107 | 1917 | 0.2863 | 0.1056 | 0.0002 | -0.59 | 0.75 | 1.34 |
| 18046 | Nfyc | 1481 | 1167 | 1329 | 0.1171 | 0.5806 | 0.0033 | 0.43 | -0.17 | -0.60 |
| 22259 | Nr1h3 | 1616 | 905 | 1084 | 0.6795 | 0.1338 | 0.0108 | 0.44 | -1.13 | -1.56 |
| 22026 | Nr2c2 | 883 | 546 | 769 | 0.0126 | 0.8108 | 0.0004 | 1.03 | -0.14 | -1.17 |
| 15370 | Nr4a1 | 68431 | 27561 | 52415 | 0.0278 | 0.4582 | 0.0001 | 1.54 | -0.59 | -2.13 |
| 18227 | Nr4a2 | 2259 | 1093 | 1327 | 0.7947 | 0.1049 | 0.0153 | 0.42 | -1.72 | -2.14 |
| 211323 | Nrg1 | 611 | 30 | 447 | 0.0000 | 0.0781 | 0.0000 | 4.42 | -1.88 | -6.31 |
| 11545 | Parp1 | 6795 | 6616 | 4924 | 0.0000 | 0.0002 | 0.4446 | -0.99 | -0.79 | 0.20 |
| 18514 | Pbx1 | 494 | 74 | 392 | 0.0000 | 0.8843 | 0.0000 | 3.74 | 0.17 | -3.56 |
| 71041 | Pcgf6 | 214 | 303 | 262 | 0.1731 | 0.6214 | 0.0093 | -0.74 | 0.29 | 1.03 |
| 18673 | Phb | 1346 | 1765 | 1623 | 0.3418 | 0.3749 | 0.0087 | -0.40 | 0.34 | 0.74 |
| 72057 | Phf10 | 1717 | 1200 | 1387 | 0.2828 | 0.1659 | 0.0006 | 0.43 | -0.48 | -0.91 |
| 54711 | Plagl2 | 1673 | 1064 | 1318 | 0.3270 | 0.3140 | 0.0048 | 0.60 | -0.55 | -1.15 |
| 22038 | Plscr1 | 2267 | 1568 | 1252 | 0.3298 | 0.0000 | 0.0041 | -0.53 | -1.56 | -1.03 |
| 18987 | Pou2f2 | 1579 | 1174 | 1694 | 0.0253 | 0.3548 | 0.1904 | 1.37 | 0.63 | -0.75 |
| 110593 | Prdm2 | 1343 | 930 | 1165 | 0.1917 | 0.7206 | 0.0189 | 0.70 | -0.22 | -0.92 |
| 50907 | Preb | 5541 | 4550 | 5409 | 0.0045 | 0.7332 | 0.0057 | 0.62 | 0.10 | -0.51 |
| 19401 | Rara | 6451 | 1227 | 4239 | 0.0000 | 0.0000 | 0.0000 | 2.11 | -2.25 | -4.36 |
| 19411 | Rarg | 1636 | 534 | 1179 | 0.0016 | 0.0917 | 0.0000 | 1.70 | -0.96 | -2.66 |
| 19664 | Rbpj | 2661 | 2370 | 1877 | 0.0836 | 0.0168 | 0.7681 | -0.72 | -0.86 | -0.14 |
| 19696 | Rel | 3643 | 1951 | 2481 | 0.5701 | 0.1637 | 0.0072 | 0.59 | -1.09 | -1.67 |
| 19697 | Rela | 13983 | 8797 | 12152 | 0.0966 | 0.8398 | 0.0121 | 0.97 | -0.16 | -1.13 |
| 19712 | Rest | 3506 | 2201 | 2713 | 0.2720 | 0.1897 | 0.0008 | 0.58 | -0.61 | -1.19 |
| 19725 | Rfx2 | 497 | 71 | 319 | 0.0010 | 0.0000 | 0.0000 | 2.32 | -3.51 | -5.83 |
| 19726 | Rfx3 | 143 | 67 | 114 | 0.0162 | 0.5397 | 0.0001 | 1.37 | -0.43 | -1.80 |
| 67150 | Rnf141 | 390 | 306 | 256 | 0.3469 | 0.0052 | 0.1425 | -0.49 | -1.06 | -0.57 |
| 19885 | Rorc | 79 | 147 | 175 | 0.3069 | 0.0347 | 0.4618 | 1.45 | 2.33 | 0.89 |
| 12394 | Runx1 | 6076 | 3864 | 4642 | 0.2703 | 0.0927 | 0.0001 | 0.51 | -0.65 | -1.16 |
| 12393 | Runx2 | 775 | 374 | 495 | 0.3856 | 0.0066 | 0.0000 | 0.62 | -1.40 | -2.02 |
| 66118 | Sarnp | 1783 | 1246 | 1473 | 0.2427 | 0.3076 | 0.0020 | 0.50 | -0.40 | -0.90 |
| 20230 | Satb1 | 110 | 36 | 88 | 0.0124 | 0.8422 | 0.0006 | 2.16 | -0.26 | -2.41 |
| 17127 | Smad3 | 1467 | 1085 | 1439 | 0.0123 | 0.6555 | 0.0248 | 0.98 | 0.22 | -0.75 |
| 17128 | Smad4 | 4589 | 3182 | 3862 | 0.0365 | 0.2682 | 0.0000 | 0.59 | -0.33 | -0.92 |
| 20665 | Sox10 | 433 | 908 | 1080 | 0.1615 | 0.0048 | 0.3372 | 1.71 | 2.74 | 1.03 |
| 109032 | Sp110 | 2682 | 1658 | 2251 | 0.0000 | 0.2612 | 0.0000 | 0.88 | -0.29 | -1.17 |
| 78912 | Sp2 | 2401 | 1543 | 1839 | 0.1993 | 0.0418 | 0.0000 | 0.49 | -0.64 | -1.13 |
| 20687 | Sp3 | 3293 | 2383 | 2852 | 0.3351 | 0.6798 | 0.0468 | 0.56 | -0.25 | -0.81 |
| 30051 | Spdef | 56 | 123 | 133 | 0.6395 | 0.0474 | 0.1785 | 0.71 | 2.04 | 1.33 |
| 20375 | Spi1 | 34580 | 29061 | 34728 | 0.0359 | 0.6081 | 0.0889 | 0.65 | 0.19 | -0.46 |
| 20728 | Spic | 219 | 304 | 397 | 0.0113 | 0.0007 | 0.7002 | 1.86 | 2.20 | 0.34 |
| 20788 | Srebf2 | 7242 | 4204 | 5269 | 0.2875 | 0.0671 | 0.0001 | 0.59 | -0.82 | -1.41 |
| 20843 | Stag2 | 1993 | 1469 | 1960 | 0.0113 | 0.6417 | 0.0247 | 0.99 | 0.23 | -0.76 |
| 20847 | Stat2 | 6565 | 4132 | 5575 | 0.0291 | 0.6037 | 0.0003 | 0.88 | -0.25 | -1.13 |
| 20848 | Stat3 | 27528 | 13577 | 21046 | 0.0003 | 0.0526 | 0.0000 | 1.11 | -0.63 | -1.73 |
| 20849 | Stat4 | 430 | 202 | 248 | 0.6937 | 0.0102 | 0.0002 | 0.44 | -1.82 | -2.25 |
| 20850 | Stat5a | 2513 | 1500 | 2086 | 0.0896 | 0.6209 | 0.0026 | 0.94 | -0.32 | -1.25 |
| 20851 | Stat5b | 6105 | 3093 | 5117 | 0.0000 | 0.5122 | 0.0000 | 1.38 | -0.20 | -1.59 |
| 20852 | Stat6 | 19498 | 14097 | 18237 | 0.0114 | 0.9386 | 0.0039 | 0.85 | 0.04 | -0.81 |
| 20872 | Stk16 | 1394 | 1089 | 1407 | 0.0005 | 0.3653 | 0.0074 | 0.92 | 0.29 | -0.63 |
| 20024 | Sub1 | 10525 | 6461 | 9320 | 0.0076 | 0.9219 | 0.0006 | 1.10 | -0.07 | -1.16 |
| 237336 | Tbpl1 | 2107 | 1945 | 2262 | 0.0032 | 0.1074 | 0.2079 | 0.59 | 0.34 | -0.25 |
| 21781 | Tfdp1 | 3032 | 3064 | 2337 | 0.0000 | 0.0007 | 0.1578 | -0.95 | -0.65 | 0.29 |
| 21426 | Tfec | 443 | 282 | 424 | 0.0370 | 0.7664 | 0.0452 | 1.32 | 0.25 | -1.07 |
| 59016 | Thap11 | 2012 | 1640 | 2135 | 0.0055 | 0.2606 | 0.1037 | 0.98 | 0.45 | -0.54 |
| 21833 | Thra | 154 | 124 | 163 | 0.0306 | 0.4030 | 0.1823 | 1.03 | 0.45 | -0.58 |
| 21807 | Tsc22d1 | 2034 | 1011 | 1829 | 0.0579 | 0.8870 | 0.0399 | 1.75 | 0.19 | -1.56 |
| 68842 | Tulp4 | 473 | 680 | 582 | 0.1453 | 0.6395 | 0.0074 | -0.81 | 0.29 | 1.10 |
| 22278 | Usf1 | 3170 | 2810 | 3711 | 0.0002 | 0.0130 | 0.2502 | 1.15 | 0.78 | -0.37 |
| 22344 | Vezf1 | 1299 | 1021 | 1292 | 0.0011 | 0.5034 | 0.0060 | 0.83 | 0.22 | -0.62 |
| 22433 | Xbp1 | 24367 | 21017 | 28799 | 0.0000 | 0.0008 | 0.1045 | 1.32 | 0.88 | -0.44 |
| 56449 | Ybx3 | 8247 | 5682 | 5151 | 0.6737 | 0.0003 | 0.0018 | -0.24 | -1.25 | -1.00 |
| 58206 | Zbtb32 | 607 | 169 | 347 | 0.4531 | 0.0101 | 0.0000 | 1.21 | -2.90 | -4.11 |
| 75580 | Zbtb4 | 159 | 313 | 259 | 0.0176 | 0.2758 | 0.0000 | -1.37 | 0.67 | 2.04 |
| 382639 | Zbtb42 | 498 | 309 | 359 | 0.5696 | 0.1097 | 0.0034 | 0.40 | -0.84 | -1.24 |
| 16969 | Zbtb7a | 5638 | 4535 | 5402 | 0.0037 | 0.8790 | 0.0016 | 0.61 | 0.05 | -0.56 |
| 24136 | Zeb2 | 7645 | 5113 | 6252 | 0.2078 | 0.3455 | 0.0019 | 0.59 | -0.42 | -1.01 |
| 21769 | Zfand3 | 648 | 716 | 772 | 0.2206 | 0.0258 | 0.5358 | 0.35 | 0.51 | 0.16 |
| 11906 | Zfhx3 | 851 | 1509 | 1169 | 0.0006 | 0.5968 | 0.0000 | -1.70 | 0.35 | 2.05 |
| 22640 | Zfp1 | 227 | 292 | 268 | 0.3299 | 0.4477 | 0.0129 | -0.41 | 0.29 | 0.70 |
| 63872 | Zfp296 | 184 | 75 | 151 | 0.0892 | 0.8879 | 0.0144 | 1.77 | -0.21 | -1.98 |
| 22695 | Zfp36 | 138207 | 77852 | 88657 | 0.6789 | 0.0076 | 0.0001 | 0.31 | -1.27 | -1.59 |
| 22751 | Zfp90 | 457 | 894 | 782 | 0.0386 | 0.0601 | 0.0000 | -0.98 | 0.84 | 1.81 |
| 626848 | Zfp971 | 554 | 86 | 402 | 0.0000 | 0.0229 | 0.0000 | 2.89 | -1.18 | -4.07 |
| 22761 | Zfpm1 | 355 | 490 | 382 | 0.0105 | 0.9862 | 0.0023 | -1.22 | -0.02 | 1.20 |
| 22697 | Zscan21 | 718 | 907 | 867 | 0.6720 | 0.2937 | 0.0433 | -0.20 | 0.38 | 0.58 |

**Table S2.**
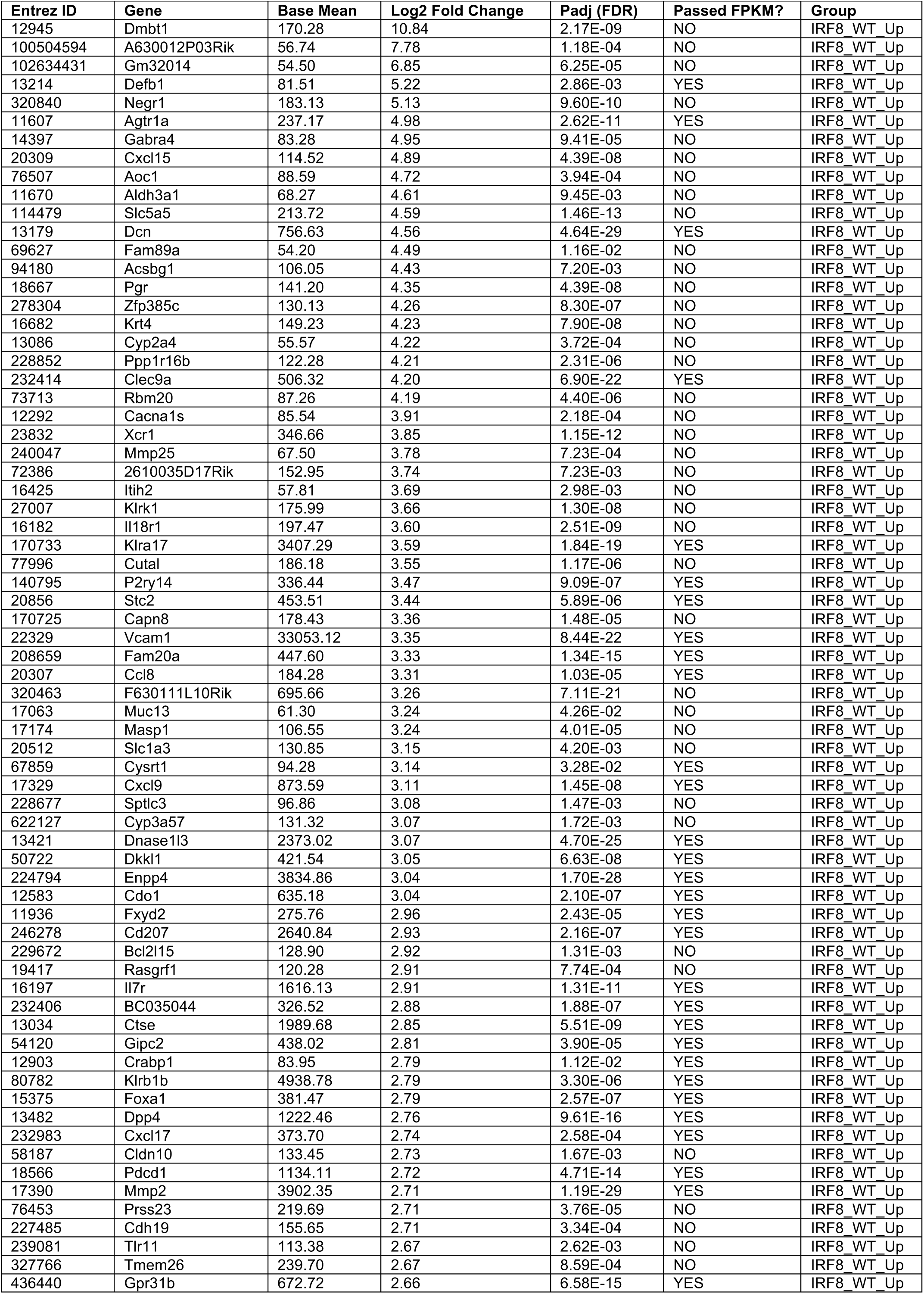
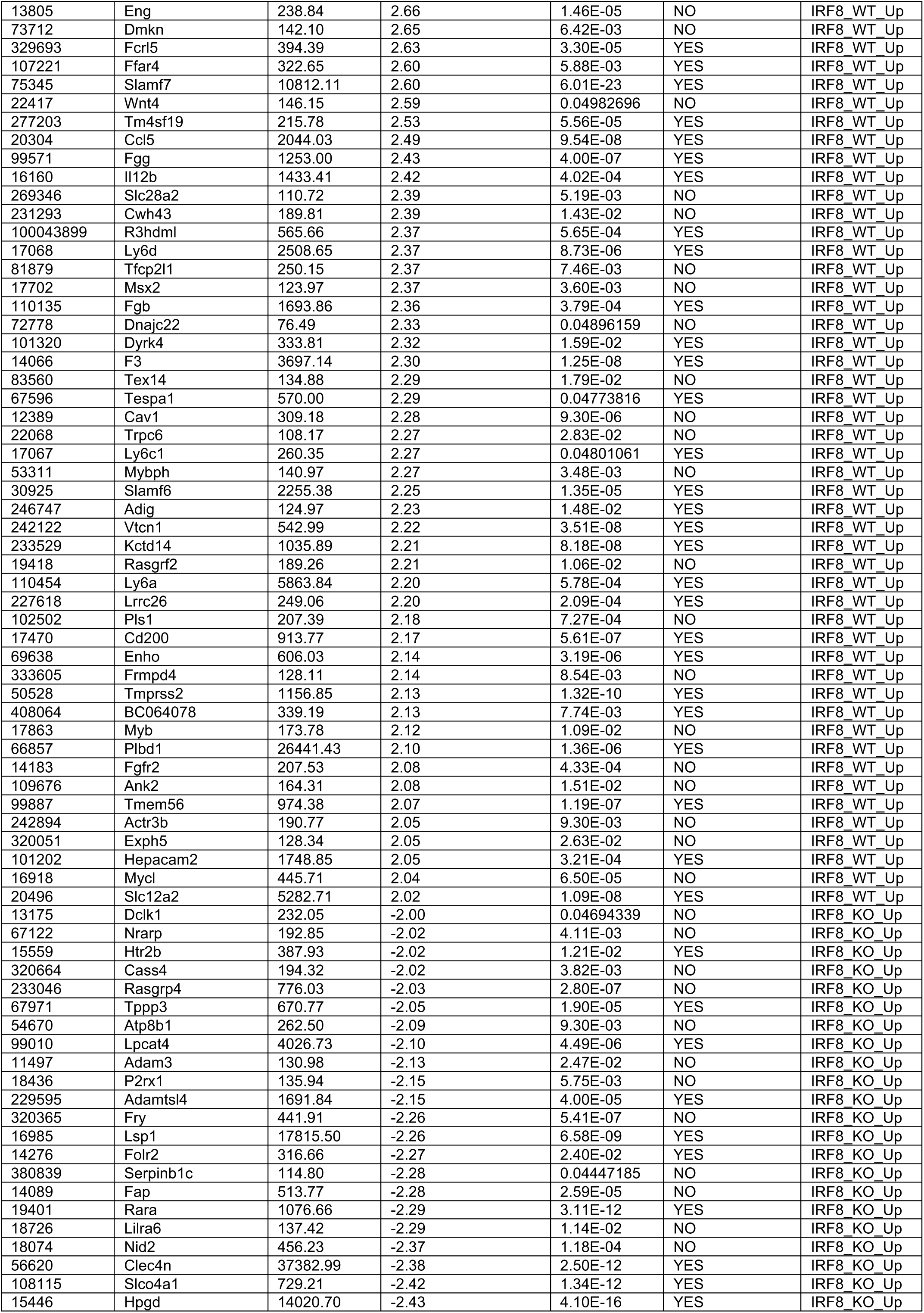
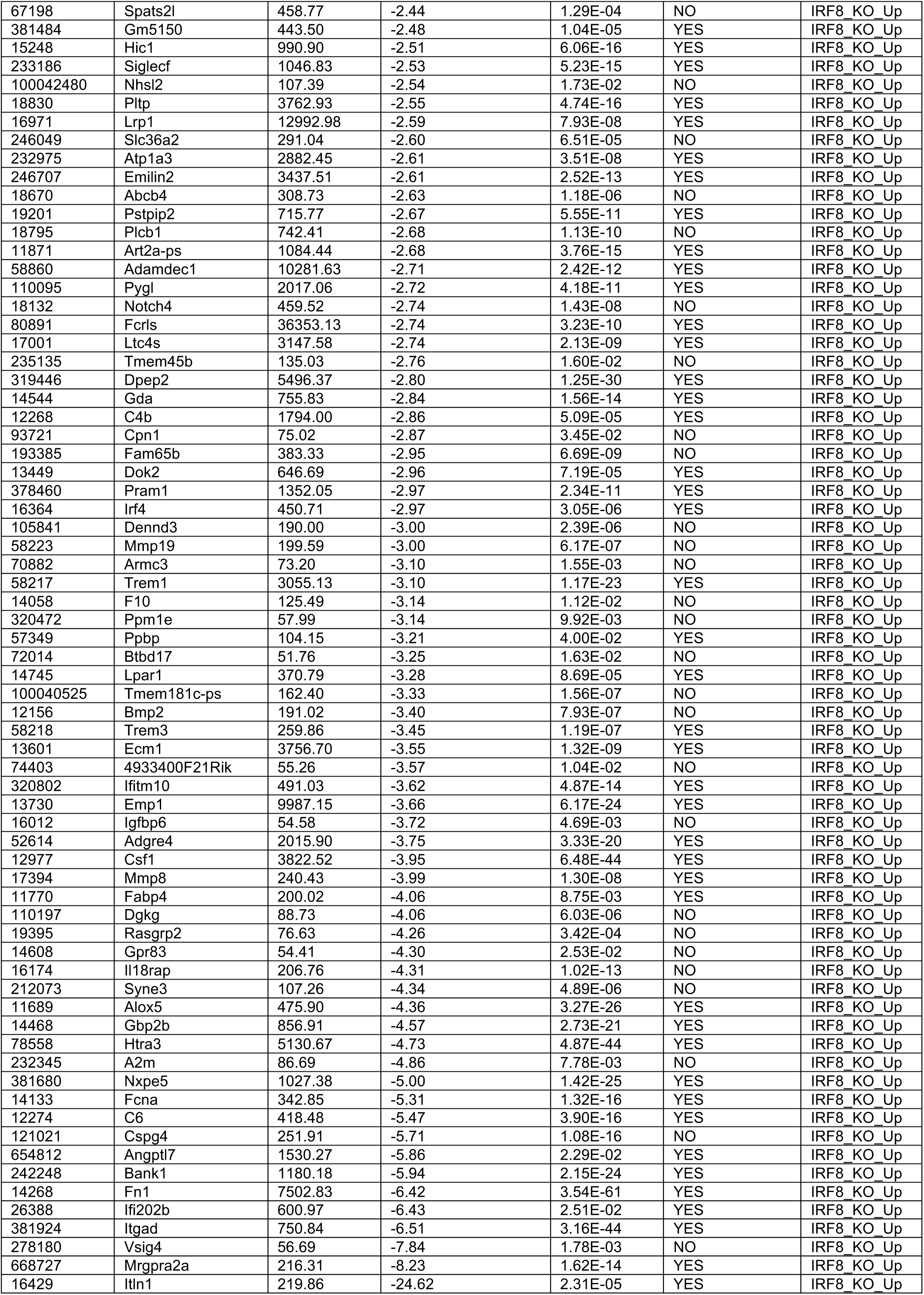
Differentially expressed genes between wild-type (WT) and IRF8-knockout (IRF8KO) TAMs.

**Table S3.**
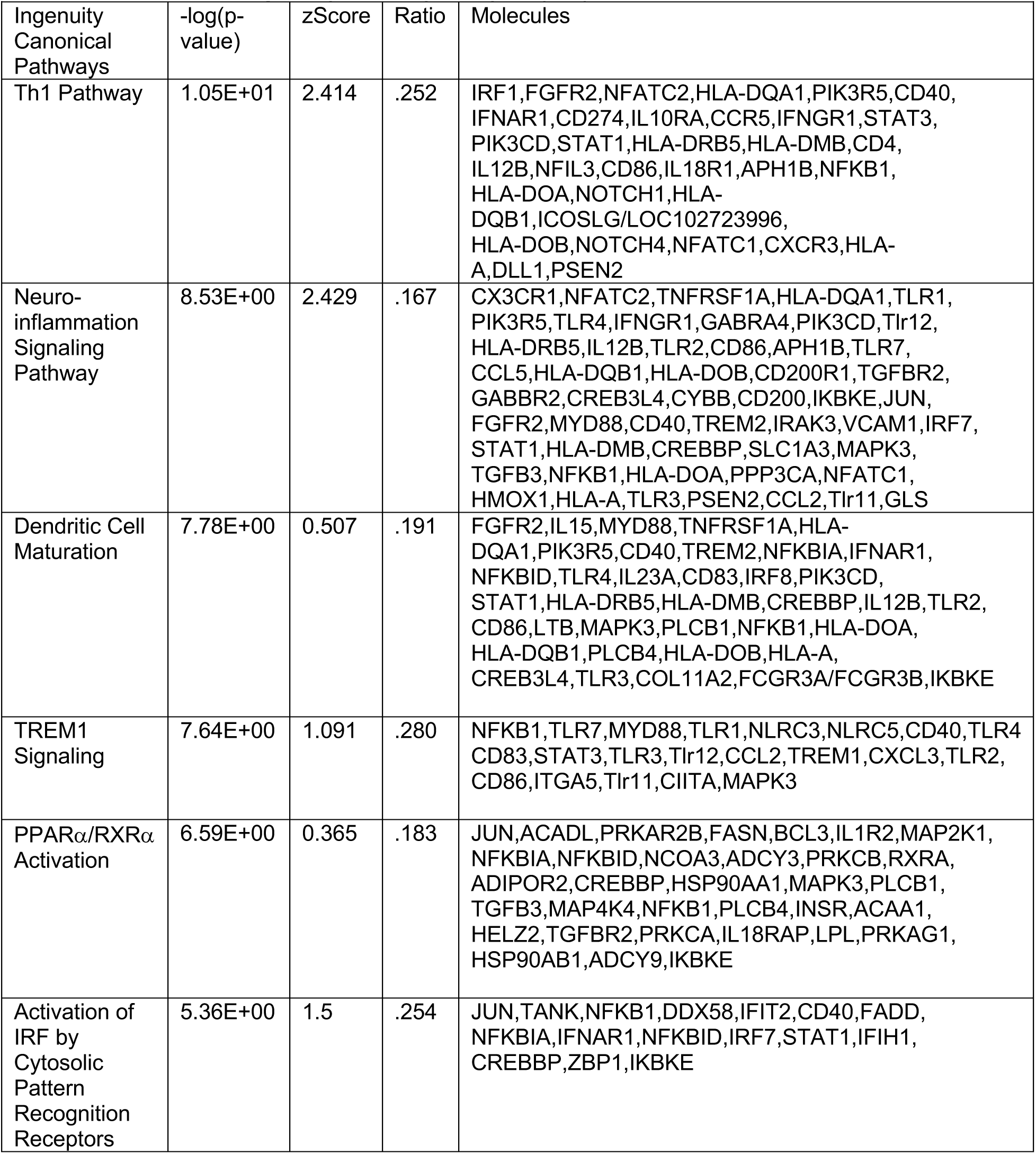
Ingenuity Pathway Analysis (IPA) for WT versus IRF8KO TAMs.

**Table S4.**
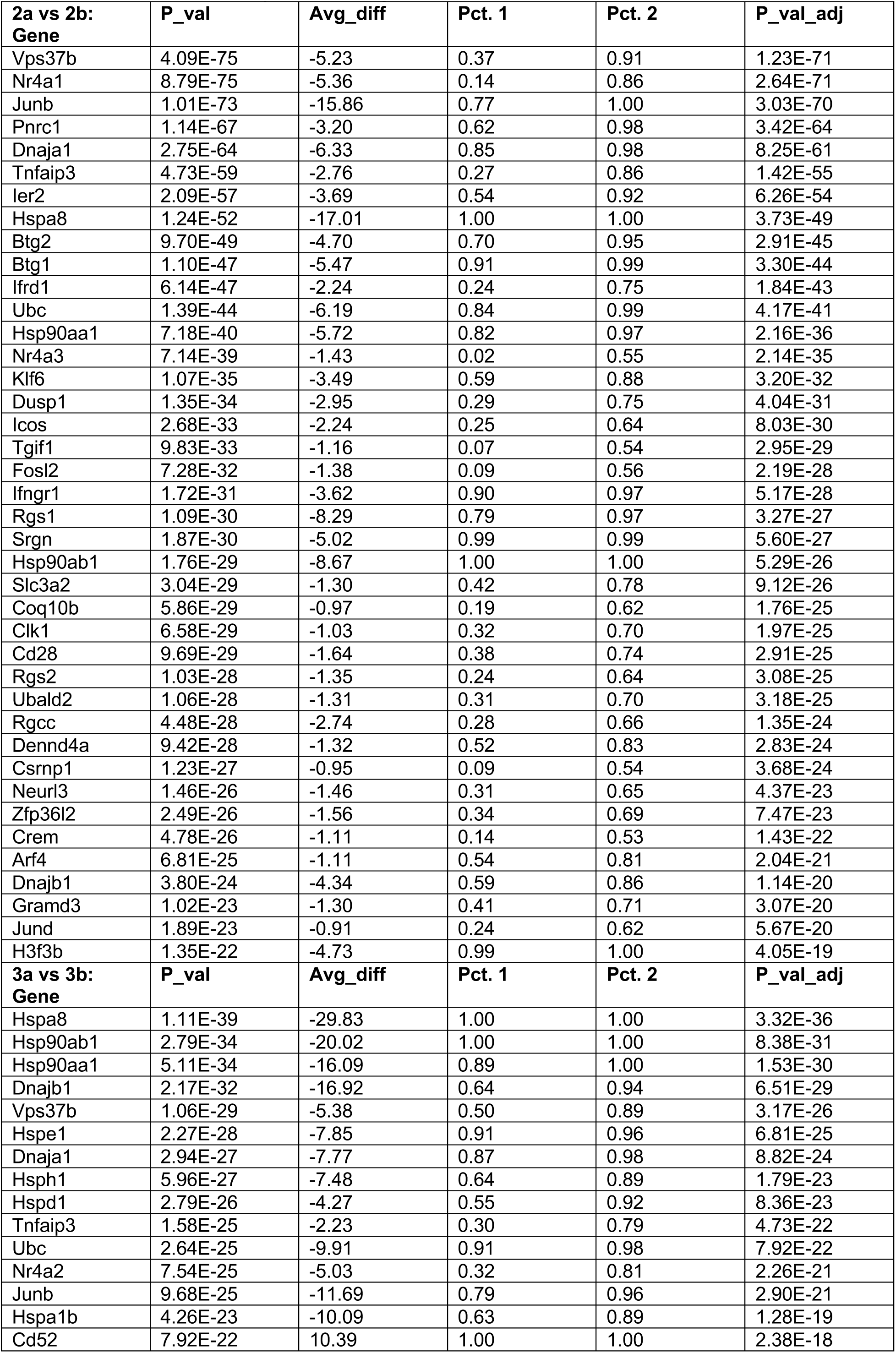
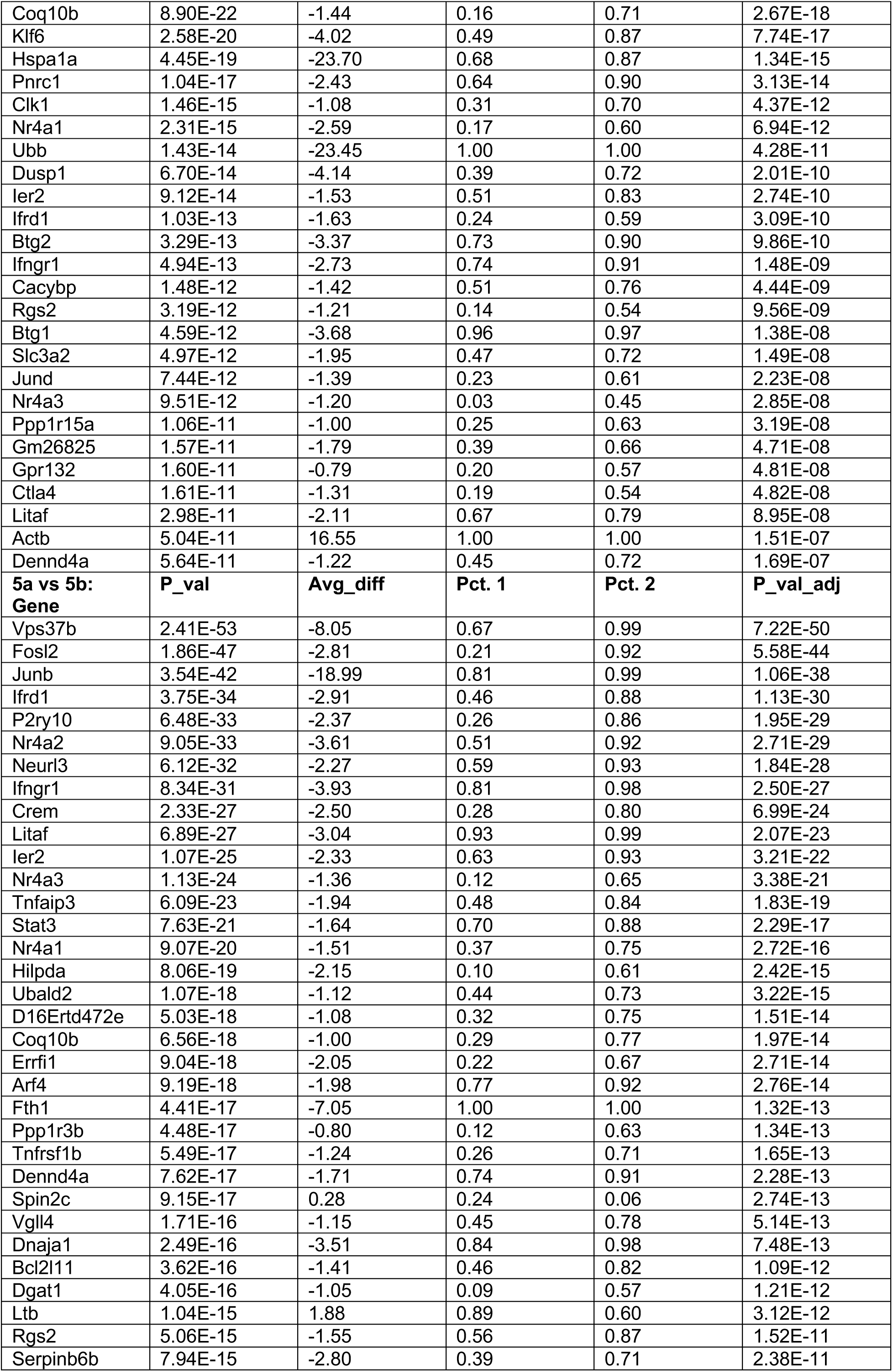
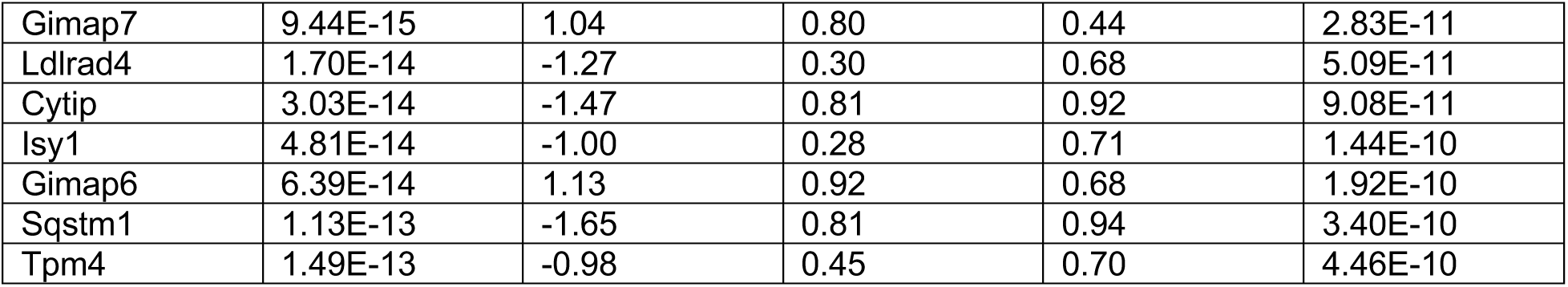
Differentially expressed genes between “a” and “b” clusters in scRNAseq.

